# Deep immune profiling reveals targetable mechanisms of immune evasion in checkpoint blockade-refractory glioblastoma

**DOI:** 10.1101/2020.12.01.404939

**Authors:** Erin F. Simonds, Edbert D. Lu, Eric V. Liu, Whitney Tamaki, Chiara Rancan, Jacob Stultz, Meenal Sinha, Lauren K. McHenry, Nicole M. Nasholm, Pavlina Chuntova, Anders Sundström, Vassilis Genoud, Shilpa A. Shahani, Leo D. Wang, Christine E. Brown, Paul R. Walker, Fredrik J. Swartling, Lawrence Fong, Hideho Okada, William A. Weiss, Mats Hellström

**Affiliations:** Department of Neurology, University of California, San Francisco, CA, USA; Division of Hematology/Oncology, Department of Medicine, San Francisco, CA, USA; Department of Neurological Surgery, University of California, San Francisco, CA, USA; Department of Immunology, Genetics and Pathology, Uppsala University, Uppsala, Sweden; Translational Research Centre in Oncohaematology, Faculty of Medicine, University of Geneva, Geneva, Switzerland; Department of Pediatrics, City of Hope, Duarte, CA, USA; Department of Pediatrics, University of California, San Francisco, CA, USA; Helen Diller Family Comprehensive Cancer Center, University of California, San Francisco, CA, USA

## Abstract

**Background:** Glioblastoma (GBM) is refractory to checkpoint blockade immunotherapy (CPI). We sought to determine to what extent this immune evasion is due to intrinsic properties of the tumor cells versus the specialized immune context of the brain, and if it can be reversed.

**Methods:** We used CyTOF mass cytometry to compare the tumor immune microenvironments (TIME) of human tumors that are generally CPI-refractory (GBM and sarcoma) or CPI-responsive (renal cell carcinoma), as well as mouse models of GBM that are CPI-responsive (GL261) or CPI-refractory (SB28). We further compared SB28 tumors grown intracerebrally versus subcutaneously to determine how tumor site affects TIME and responsiveness to dual CTLA-4/PD-1 blockade. Informed by these data, we explored rational immunotherapeutic combinations.

**Results:** CPI-sensitivity in human and mouse tumors was associated with increased T cells and dendritic cells, and fewer myeloid cells, in particular PD-L1+ tumor associated macrophages. The SB28 mouse model of GBM responded to CPI when grown subcutaneously but not intracerebrally, providing a system to explore mechanisms underlying CPI resistance in GBM. The response to CPI in the subcutaneous SB28 model required CD4 T cells and NK cells, but not CD8 T cells. Recombinant FLT3L expanded dendritic cells, improved antigen-specific T cell priming, and prolonged survival of mice with intracerebral SB28 tumors, but at the cost of increased Tregs. Targeting PD-L1 also prolonged survival, especially when combined with stereotactic radiation.

**Conclusions:** Our data suggest that a major obstacle for effective immunotherapy of GBM is the low antigenicity of the tumor cells coupled with poor antigen presentation in the brain, rather than intrinsic immunosuppressive properties of GBM tumor cells. Deep immune profiling identified dendritic cells and PD-L1+ tumor-associated macrophages as promising targetable cell populations, which was confirmed using therapeutic interventions *in vivo*.

**Graphical Abstract:** 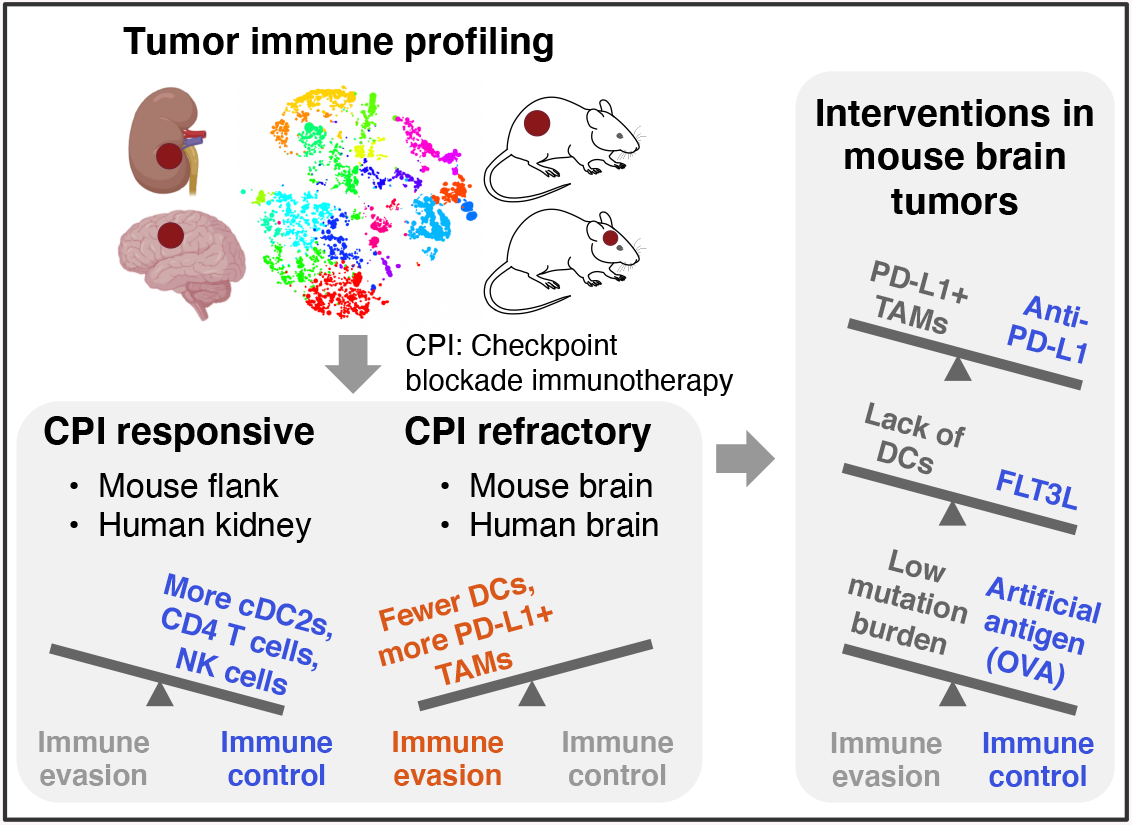

*In Brief:* In mice and humans, tumors that were sensitive to checkpoint blockade had consistent immunological features. A mouse model of glioma that is refractory to checkpoint blockade was sensitized by increasing antigen presentation through a variety of approaches.

## BACKGROUND

Glioblastoma (GBM) is the most common primary malignant brain tumor. Despite progress understanding the genetics and biology of GBM, the prognosis remains dismal and treatment options are limited beyond maximal safe resection, radio-chemotherapy and Tumor Treating Fields, resulting in a median survival of less than two years from diagnosis^1^.

Stimulation of the adaptive immune response using checkpoint blockade immunotherapy (CPI) has revolutionized treatment for melanoma, lung, bladder, and renal cancers^2^. The clinically approved CPI drugs target two classes of lymphocyte inhibitory pathways (CTLA-4, PD-1/PD-L1), which are particularly important in regulating priming and activation of T cells. Several clinical trials have evaluated the efficacy of CPI in glioblastoma, but outcomes have been generally negative^3,4^, even in somatically hypermutated GBM^5^.

The lack of clinically meaningful responses to CPI in GBM has been attributed to the specialized immune environment of the brain, which includes mechanical obstruction at the blood-brain-barrier, unique brain-resident phagocytes (microglia and border-associated macrophages), unconventional lymphatic drainage, and a generally immunosuppressive environment^6,7^. However, the clinical failures of dual CTLA-4/PD-1 blockade in GBM cannot be entirely attributed to the immune specialization of the brain, as brain metastases from melanoma and non-small-cell lung cancer are CPI-responsive^8^. Why, then, do gliomas resist immune checkpoint blockade, while brain metastases from melanomas and carcinomas generally respond?

Recently, two groups performed single-cell profiling of the TIME of primary brain malignancies and brain metastases to explore whether tumor origin influences immune cell recruitment^9,10^. They showed that the TIME was shaped by the origin of the primary tumor – gliomas and ependymomas had reduced lymphocyte infiltration and more tumor-associated macrophages (TAMs), compared to melanoma or carcinoma brain metastases. TAMs were also skewed phenotypically based on their ontogeny as tissue-resident microglia or blood-derived monocytic macrophages, as well as the origin of the primary tumor. Brain tumors, both primary and metastatic had uniformly poor infiltration of dendritic cells (DCs). These immune profiling efforts suggest that the myeloid cells in brain tumors are shaped by origin of the primary tumor, and that they can support a productive immune response in the context of carcinoma metastases, but not in GBM.

Thus, evidence in humans suggests that the hurdles of effective immunotherapy of GBM are at least two-fold. Tumor-intrinsic properties affect the quantity and quality of the TIME. However, the extent to which tumor-extrinsic properties of the brain – unique cell types and poor antigen drainage to lymph nodes – influence T cell priming has not been addressed in the context of response to immune checkpoint blockade. Here, we combine mass cytometry profiling of human and mouse tumors to reveal that the makeup of the TIME is influenced by the origin of the primary tumor, as well as the site of growth. Furthermore, we show that certain mechanisms mediating intracerebral immune evasion can be targeted to improve survival.

## METHODS

Please refer to the Online Supplemental Methods.

## RESULTS

### CPI-refractory GBM is associated with abundant PD-L1+ tumor-associated macrophages and poor T cell infiltration

We compared the immune cell populations in two CPI-refractory tumor types, GBM and sarcoma, to renal cell carcinoma (RCC), which is often CPI-responsive. Single-cell profiling of 19 GBM, 11 RCC, and 4 sarcoma tumors was performed using mass cytometry^11^ (**Figure 1A**, **Suppl Table S1**). To improve the unsupervised detection of immune cell subsets, we included samples of healthy PBMC (PHA-stimulated and non-stimulated) from one normal donor, as well as paired normal tissue from one RCC patient and one sarcoma patient. Samples were stained with two antibody panels focusing on either T cells or myeloid cells, each measuring 42 markers (**Suppl Table S2 & S3**). Single-cell data was processed with the FlowSOM^12^ and PhenoGraph^13^ clustering algorithms to reduce the complexity from millions of CD45-positive cells to 22 metaclusters representing phenotypically distinct immune cell subsets (details in **Materials and Methods**). Immune subset (e.g., cluster) abundances from the T cell-focused staining panel were visualized as a reduced-dimensionality map using the *t*-SNE algorithm^14,15^. In this high-level view of TIME landscapes across the three tumor types, all GBM samples were clearly separated from RCC and sarcoma, which were intermingled (**Figure 1A** and **Suppl Figure S1A,B**). To explore this in more detail, we directly compared the immune cell subset abundances in GBM versus RCC using the edgeR algorithm^16^. We found that nearly all subsets of T cells, plus NK cells, were decreased in the GBM TIME (**Figure 1B**), consistent with prior reports^17^. Using a second, myeloid-focused staining panel to compare GBM to RCC, we observed an increase in multiple TAM subsets in GBM, including a large population of PD-L1+ CD68+ TAMs (**Figure 1C**). This is consistent with a report that a PD-L1+ myeloid cell subset was specific to GBM as compared to renal cell, colorectal, prostate and non-small cell lung cancer^17^.

**Figure 1.**
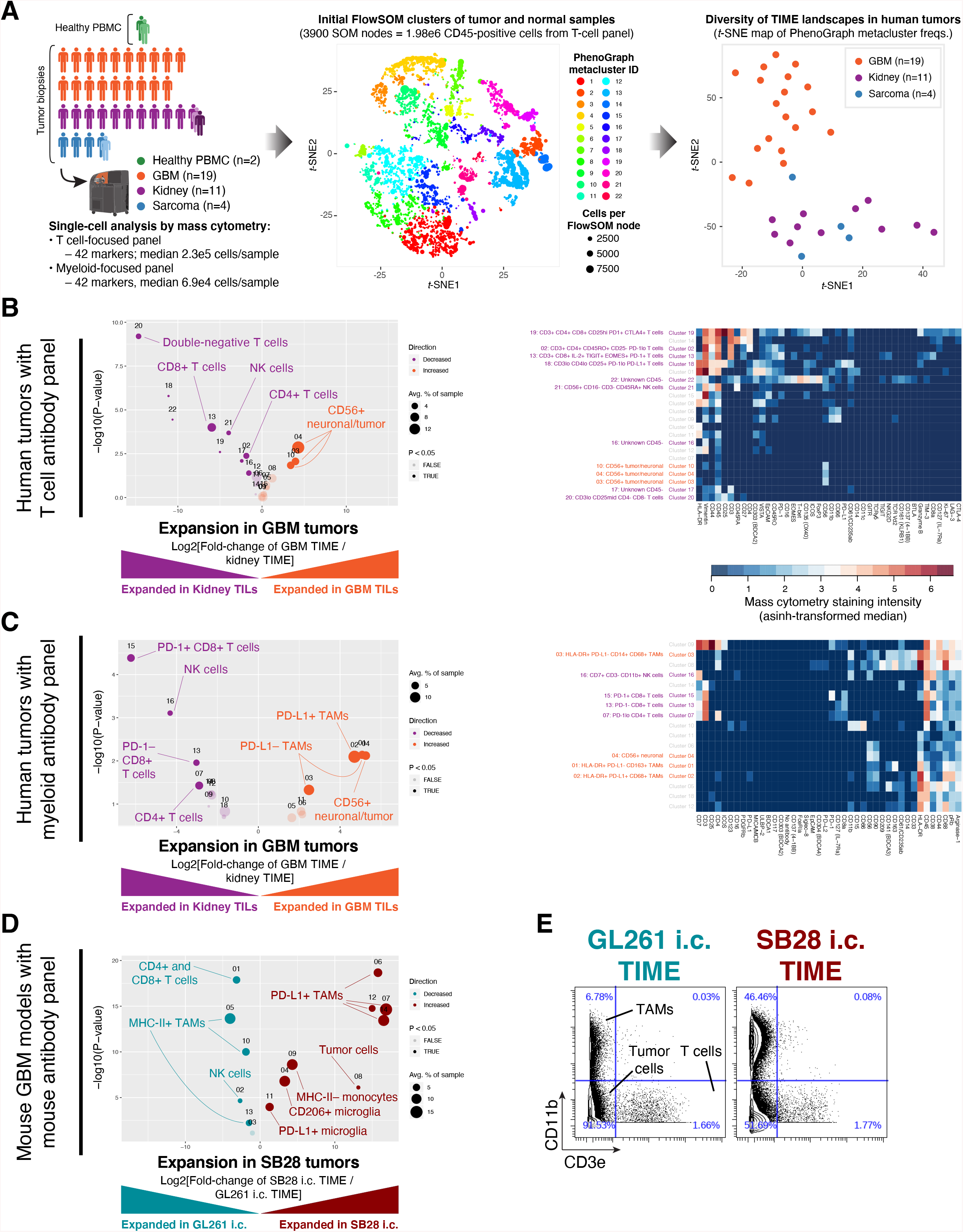
Abundant PD-L1+ tumor-associated macrophages and lack of MHC-II+ antigen-presenting cells are associated with resistance to dual CTLA-4/PD-1 checkpoint blockade. **A**. Immune profiles were obtained from 39 samples: 19 GBM primary tumor biopsies, 11 RCC primary tumor biopsies (one with paired tumor-adjacent normal tissue and metastatic lesion), 4 sarcoma primary tumor biopsies (one with paired tumor-adjacent normal tissue) and PBMCs from 1 healthy donor (with paired unstimulated and phytohemagglutinin-stimulated aliquots). Single-cell mass cytometry data were acquired using T cell-focused and myeloid-focused antibody panels with 42 markers each (see **Materials & Methods** and **Suppl Tables S2 and S3**). Data were filtered on CD45-positive cells and processed using the PhenoGraph + FlowSOM analysis pipeline to segregate immune cell types into metaclusters and quantify their frequency across samples. To compare the overall tumor immune microenvironment (TIME) landscape across patients, metacluster frequencies for each primary tumor were fed into the *t*-stochastic neighbor embedding (*t*-SNE) dimensionality reduction algorithm to produce a map of TIME landscapes, organized spatially by similarity. Data from the T cell panel is shown in all plots; corresponding plots for the myeloid panel are in **Figures S1A-B**. **B**. Volcano plot comparing abundance of immune cell populations (clusters) in GBM (orange) versus RCC (purple) tumor biopsies stained with the CyTOF human T-cell antibody panel. Statistically significant clusters in volcano plots are highlighted in opaque color and indicated with a cell type label. Diameter of the circle indicates the mean frequency of cells in the sample assigned to that cluster. Heatmap (right panel) indicates manually defined cluster phenotypes and median intensity of antibody staining in each cluster. **C**. Volcano plot and heatmap as in (B), stained with the CyTOF human myeloid antibody panel. **D**. Volcano plot as in (B) comparing abundance of immune cell populations in mouse GBM models SB28 (red) and GL261 (teal) stained with the CyTOF mouse antibody panel. **E**. Biaxial plots of representative raw CyTOF single-cell measurements of CD11b and CD3e on dissociated GL261 or SB28 tumors. These represent two of the 42 CyTOF mass cytometry channels, and two of the tumor biopsies, used to produce the volcano plot in (D). The staining patterns typical of tumor-associated macrophages (CD11b+), T-cells (CD3e+), and tumor cells (CD11b− CD3e−) are indicated, but clustering was performed using a total of 38 antibody markers (see **Suppl Table S4**).

The most commonly used mouse model of glioma, GL261, was established on the C57BL/6 background 50 years ago by chemical mutagenesis and was initially described as resembling ependymoblastoma by histology^18^. The SB28 mouse model of invasive glioma was developed in 2014 through simultaneous suppression of p53, overexpression of *Pdgfb* and hyperactive ERK signaling through *NRAS^G12D^*, also on the C57BL/6 background^19^. We analyzed bulk RNAseq datasets from *in vivo* orthotopic (intracerebral) SB28 and GL261 tumors and compared these to a reference cohort of human primary brain tumors. This cross-species analysis revealed that both SB28 and GL261 tumors were closely related to the mesenchymal subtype of GBM, and were distinct from other human primary brain tumors (**Suppl Figure S1D, S1E**). The two models differ markedly in response to CPI. GL261 responds to immunotherapies targeting a range of immune checkpoints including CTLA-4, PD-1, PD-L1, and TIM-3^20,21^. SB28, on the other hand, does not respond to dual CTLA-4/PD-1 blockade, and has nearly 50-fold fewer non-synonymous mutations than GL261^22^. Profiling the TIME of these two glioma models by mass cytometry (**Suppl Table S4**), we found that the CPI-resistant SB28 model had fewer T cells and more PD-L1+ TAMs than the CPI-responsive GL261 model (**Figure 1D**). Notably, the macrophage-rich, T cell-poor immune profile of SB28 was consistent with human GBM tumors (**Figure 1C**). In SB28 tumors, TAMs outnumbered T cells by 46:1, compared to 4:1 in GL261 tumors (**Figure 1D-E**), indicating markedly different states of immune activation.

To compare TAM infiltration and PD-L1 expression in human GBM versus a wide range of other cancers, we analyzed the TCGA database for mRNA expression of two TAM-associated genes (*CD163*; PD-L1 *[CD274]*) and one T cell-associated gene (*CD3E*) (**Suppl Figure S1F-H**). GBM stood out as having the highest CD163/CD3E ratio of the 30 tumor types analyzed, indicating GBM tumors have a uniquely TAM-dominated TIME combined with poor T cell infiltration.

Overall, profiling the immune landscapes of human and mouse tumors highlighted the scarcity of T cells and plenitude of PD-L1+ TAMs in GBM. In humans, these features distinguished GBM from CPI-responsive RCC, while in mice, these same features distinguished CPI-refractory SB28 from CPI-responsive GL261.

### The response to dual CTLA-4/PD-1 blockade in SB28 is dependent on tumor site and correlated with abundance of dendritic cells

Dual CTLA-4/PD-1 blockade was previously reported to prolong survival of GL261 intracerebral tumors, but to be ineffective for SB28^22^. We hypothesized that SB28 tumors might respond to CPI if grown outside of the immunologically specialized environment of the brain. We therefore compared the effectiveness of dual CTLA-4/PD-1 blockade (**Figure 2A**) on SB28 intracerebral tumors versus SB28 tumors grown subcutaneously in the flank. Consistent with the prior report, CPI treatment failed to slow the growth of established intracerebral SB28 tumors (**Figure 2B**). Surprisingly, the same CPI cocktail completely blocked growth of established subcutaneous SB28 tumors in the flank (**Figure 2C**).

**Figure 2.**
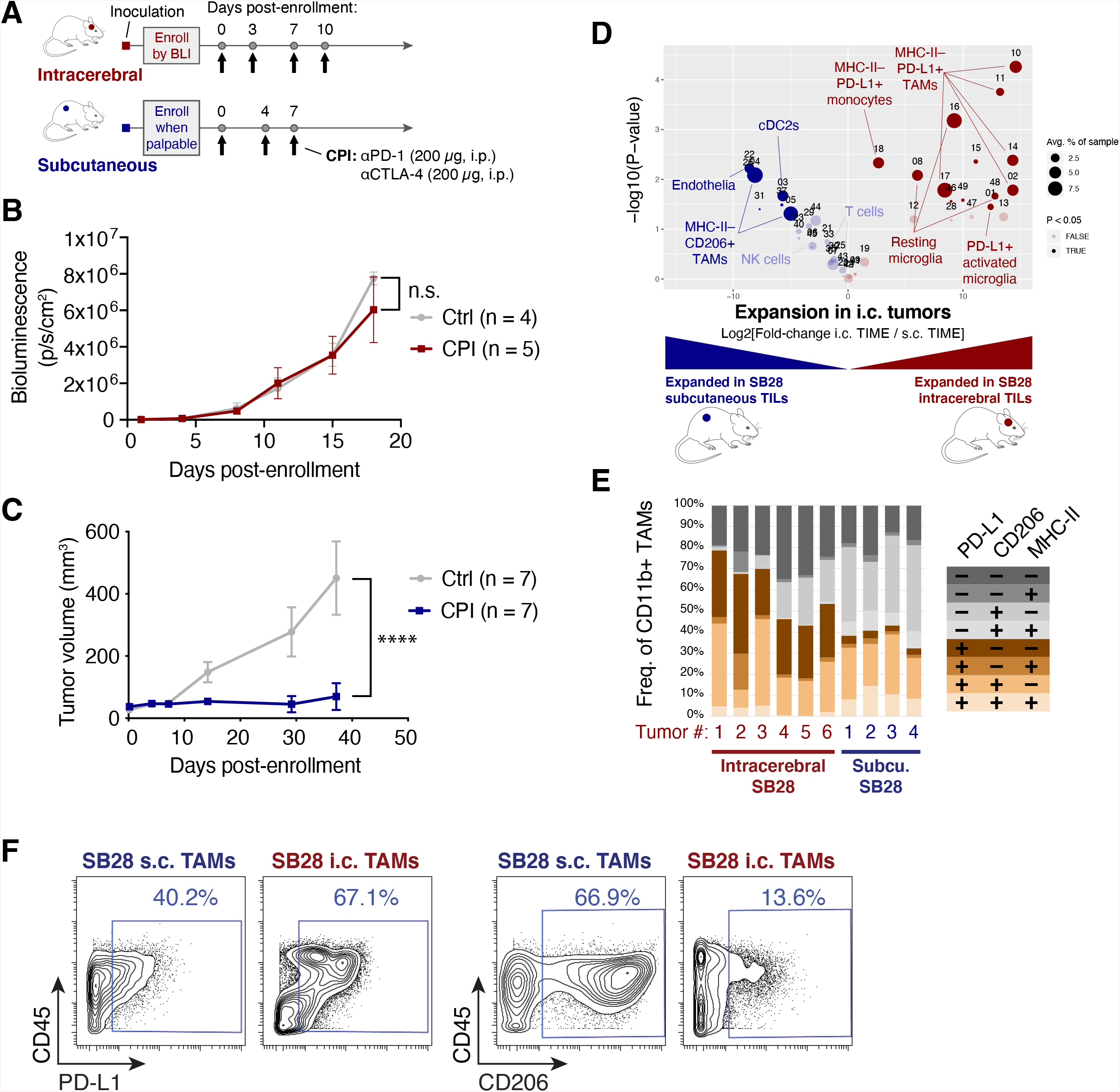
Subcutaneous SB28 tumors differ from intracerebral tumors in response to CPI and phenotype of TAMs in the TIME. **A**. Schematic illustration of dosing schedule for intracerebral or subcutaneous SB28 tumors. **B**. Bioluminescence measurements of SB28 injected intracerebrally and treated with IgG control (grey) or anti-CTLA-4 and anti-PD-1 (red). Statistical test: Mixed-effects model (n.s.: P > 0.05). Error bars: SEM. **C**. Tumor volume measurements of SB28 injected subcutaneously and treated with IgG control (grey) or anti-CTLA-4 and anti-PD-1 (blue). Statistical test: Repeated measures two-way ANOVA (****: P ≤ 0.0001). Error bars: SEM. **D**. Volcano plot comparing abundance of tumor-infiltrating leukocyte (TIL) subpopulations in dissociated intracerebral (i.c., red) and subcutaneous (s.c., blue) SB28 tumors using the CyTOF mouse immune cell panel. Statistically significant clusters in volcano plots are highlighted in opaque color and indicated with a cell type label. **E**. Mass cytometry data from SB28 subcutaneous or intracerebral tumors were manually gated on CD11b+ TAMs expressing or lacking PD-L1, CD206, or MHC-II. Frequencies of TAMs expressing all possible permutations of these three markers were quantified. **F**. Biaxial plots of representative raw CyTOF single-cell measurements of CD45, PD-L1, and CD206 on dissociated SB28 subcutaneous or intracranial tumors. Only CD11b+ events are shown. These gates were used to produce the column graph in (E).

We performed mass cytometry of dissociated untreated intracerebral and subcutaneous SB28 tumors to identify populations in the TIME that might contribute to the differential response to treatment in the brain versus the flank. Intracerebral SB28 tumors were distinguished by abundant PD-L1+ myeloid cells (TAMs and microglia) while subcutaneous SB28 tumors contained more cDC2 cells (CD11c+, CD11b+, CD103-, CD8-, Flt3^int^) which are efficient at priming CD4 T cells^23^ (**Suppl Figure S2A**). Overall, SB28 subcutaneous tumors appeared to be more inflamed and enabled for antigen presentation, due to the presence of cDC2s and a trend toward increased T cells and NK cells.

To further explore the myeloid phenotypes present in SB28 tumors grown intracerebrally or subcutaneously, we examined the co-expression of PD-L1, CD206, and MHC-II on tumor-infiltrating CD11b+ myeloid cells. The glioma TAMs in our dataset did not follow a binary split into the classic “M1” pro-inflammatory and “M2” immunosuppressive phenotypes^24^. Instead, we observed simultaneous co-expression of M1 markers (MHC-II) and M2 markers (PD-L1, CD206) in every permutation. The immune microenvironment of subcutaneous SB28 tumors was dominated by MHC-II- PD-L1- CD206+ TAMs, while intracerebral SB28 tumors were dominated by MHC-II- PD-L1+ CD206- TAMs (**Figure 2E, 2F**). The presence of MHC-II+ TAMs was a defining feature shared by SB28 subcutaneous tumors (**Figure 2E**) and GL261 intracerebral tumors (**Figure 1D**), both of which were responsive to CPI, while MHC-II+ TAMs were essentially absent in the CPI-refractory SB28 intracerebral tumors (**Figure 2E-F**).

Our data suggest that the lack of response to CPI observed in intracerebral SB28 tumors was not simply a tumor cell-intrinsic property attributable to its low neoantigen load. Rather, CPI-responsiveness could be restored by changing the anatomical tumor site, indicating that tumor cell-extrinsic mechanisms contribute to immune evasion. Single-cell profiling implicated several specific mechanisms, including antigen presentation by DCs and the balance between immunosuppressive and immune-activating TAMs.

### Dual CTLA-4/PD-1 blockade of SB28 subcutaneous tumors elicits systemic immunity, expands cDC2s and is dependent on CD4 T cells and NK cells

To investigate if the CPI-mediated tumor growth inhibition in SB28 subcutaneous tumors resulted in sustained immunological memory, we took mice cured of subcutaneous SB28 tumors and rechallenged them with intracerebral SB28 tumors (**Figure 3A**). All mice previously cured of SB28 subcutaneous tumors rejected the subsequent intracerebral SB28 tumors, while all naive mice died within 45 days of the intracerebral tumor inoculation (**Figure 3B**).

**Figure 3.**
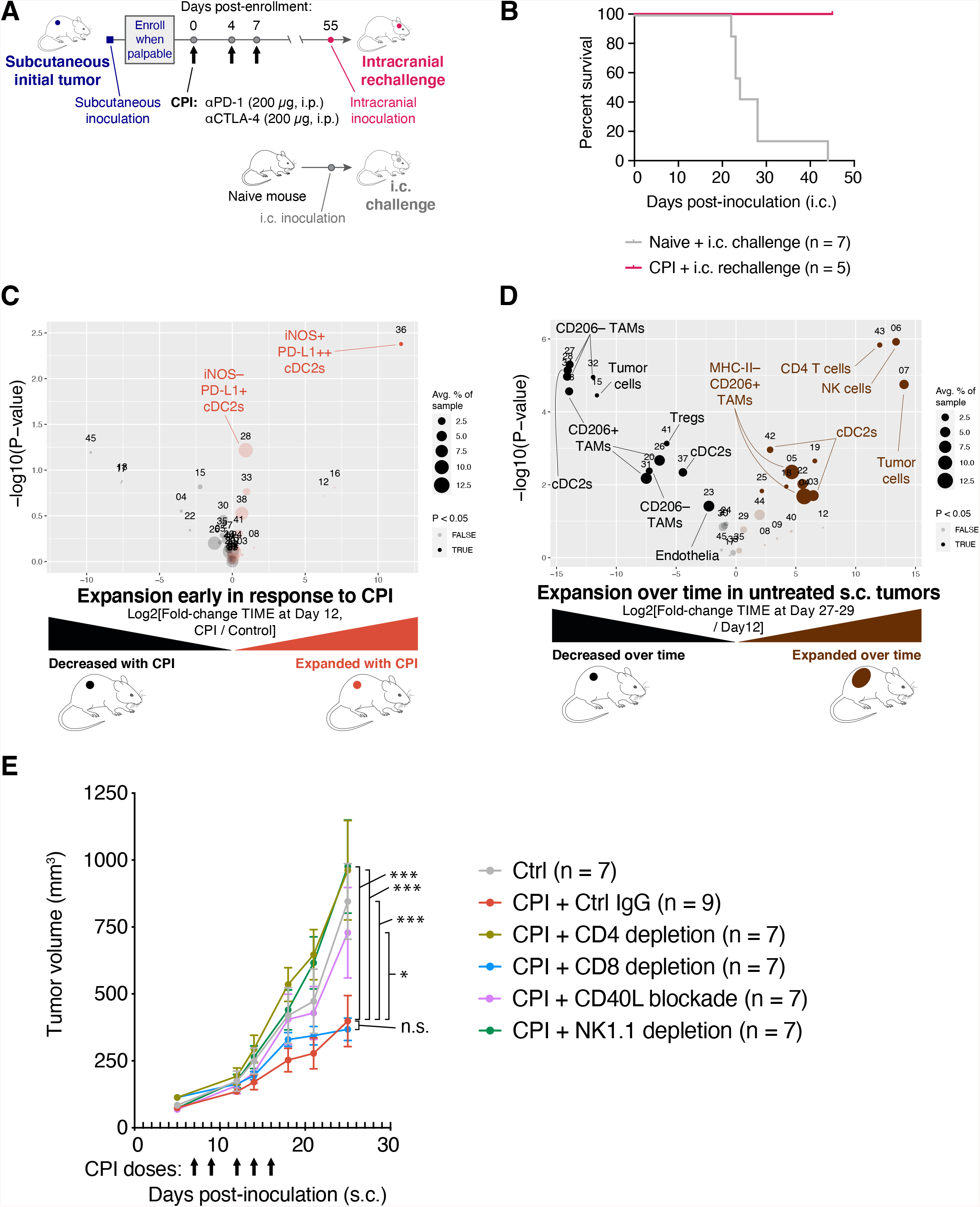
CPI treatment of SB28 subcutaneous tumors can elicit systemic memory and requires similar immune subsets to those involved in natural immune surveillance. **A**. Schematic illustration of dosing schedule for SB28 intracerebral tumor rechallenge. **B**. Mice previously cured of SB28 subcutaneous tumors (red line) were rechallenged with SB28 intracerebral tumors. Survival was compared to naive mice (grey line) challenged with SB28 intracerebral tumors and illustrated in a Kaplan-Meier curve. No treatments were administered. **C**. Volcano plot comparing abundance of immune subpopulations in dissociated subcutaneous SB28 tumors on Day 12 after CPI treatment (orange) versus isotype control treatment (black), using the mouse immune cell panel. Statistically significant clusters in volcano plots are highlighted in opaque color and indicated with a cell type label. **D**. Volcano plot as in (C) comparing abundance of immune subpopulations in dissociated subcutaneous SB28 tumors on Day 27-29 (brown) versus Day 12 (black). **E**. Tumor volume measurements of SB28 subcutaneous tumors treated with control IgG (grey) or CPI in the context of either CD4 T cell depletion (light green), CD8 T cell depletion (blue), CD40L blockade (orange), or NK cell depletion (dark green). Depleting or blocking antibodies were administered on days -3, -2 and -1 as described in **Suppl Table S6**. CPI was administered on days 7, 9, 12, 14, 16 as indicated (arrows). Statistical test: Repeated measures two-way ANOVA (***: P ≤ 0.001; *: P ≤ 0.05; n.s.: P > 0.05). Error bars: SEM.

To understand early events in the response to CPI, we performed immune profiling of subcutaneous SB28 tumors resected on day 12 of treatment with either dual CTLA-4/PD-1 or isotype control. The most prominent change was the expansion of a small iNOS+ cDC2 subset in CPI-treated subcutaneous tumors and a trend toward increased iNOS- cDC2s (**Figure 3C**).

Given that subcutaneous SB28 tumors were sufficiently immunogenic to respond to CPI, we hypothesized that they would also undergo changes over time due to endogenous immune surveillance, even in untreated mice. We compared the TIME of subcutaneous SB28 tumors at early (day 12) versus end stage (day 27-29) time points in untreated mice. The TIME changed dramatically over two weeks of unchecked tumor growth. In particular, we observed increases in subsets of cDC2s, NK cells and CD4 T cells at end stage compared to Day 12, and concomitant decreases in some subtypes of TAMs and Tregs (**Figure 3D**). Notably, NK cells were the most significantly increased subset, representing ~5% of immune cells in end stage tumors, but <0.1% of immune cells in Day 12 tumors. To identify the immune populations that mediated the anti-tumor effect of CPI in subcutaneous tumors, we performed CPI treatment concurrently with depletion of CD4 T cells, CD8 T cells, or NK cells (**Figure 3E, Suppl. Figure S3C**). We also tested a blocking antibody against CD40 ligand (CD40L; CD154) to disrupt licensing of DCs^25,26^. The efficacy of CPI in subcutaneous SB28 tumors was strongly dependent on CD4+ T cells, NK cells and CD40:CD40L interactions. There was not an absolute requirement for CD8+ T cells, but tumor regression in response to CPI was seen in 0 / 8 of the CD8-depleted mice, compared with 3 / 9 of the non-depleted mice, suggesting that CD8 T cells played a minor role in CPI-mediated tumor rejection (**Suppl Figure S3D**).

These data suggest that CPI can elicit successful priming against SB28 tumor antigens in an extracranial setting, and that tumor control in that context is a multi-step process involving NK cells, DCs, CD4 T cells, and to a lesser extent, CD8 T cells.

### Expression of a strong antigen prevents SB28 engraftment in the flank, but not in the brain

To explore tumor antigen presentation in more detail, we used SB28-OVA-FL, a variant of SB28 that was engineered to express the full-length chicken ovalbumin (OVA) gene. Full-length OVA contains at least 10 high-affinity MHC class I-restricted epitopes^27^ and several MHC class II-restricted epitopes^28^. The parental SB28 tumor line had a 100% engraftment rate at both intracerebral and subcutaneous sites while the SB28-OVA-FL line had a 38% engraftment at the intracerebral site, and a 0% engraftment rate at the subcutaneous site across multiple experiments (**Figure 4A**, **Suppl. Figure S4E**). This suggests that the intracerebral site is more permissive of tumor growth than the subcutaneous site, even when the tumor cells express multiple strong MHC class I and class II antigens.

**Figure 4.**
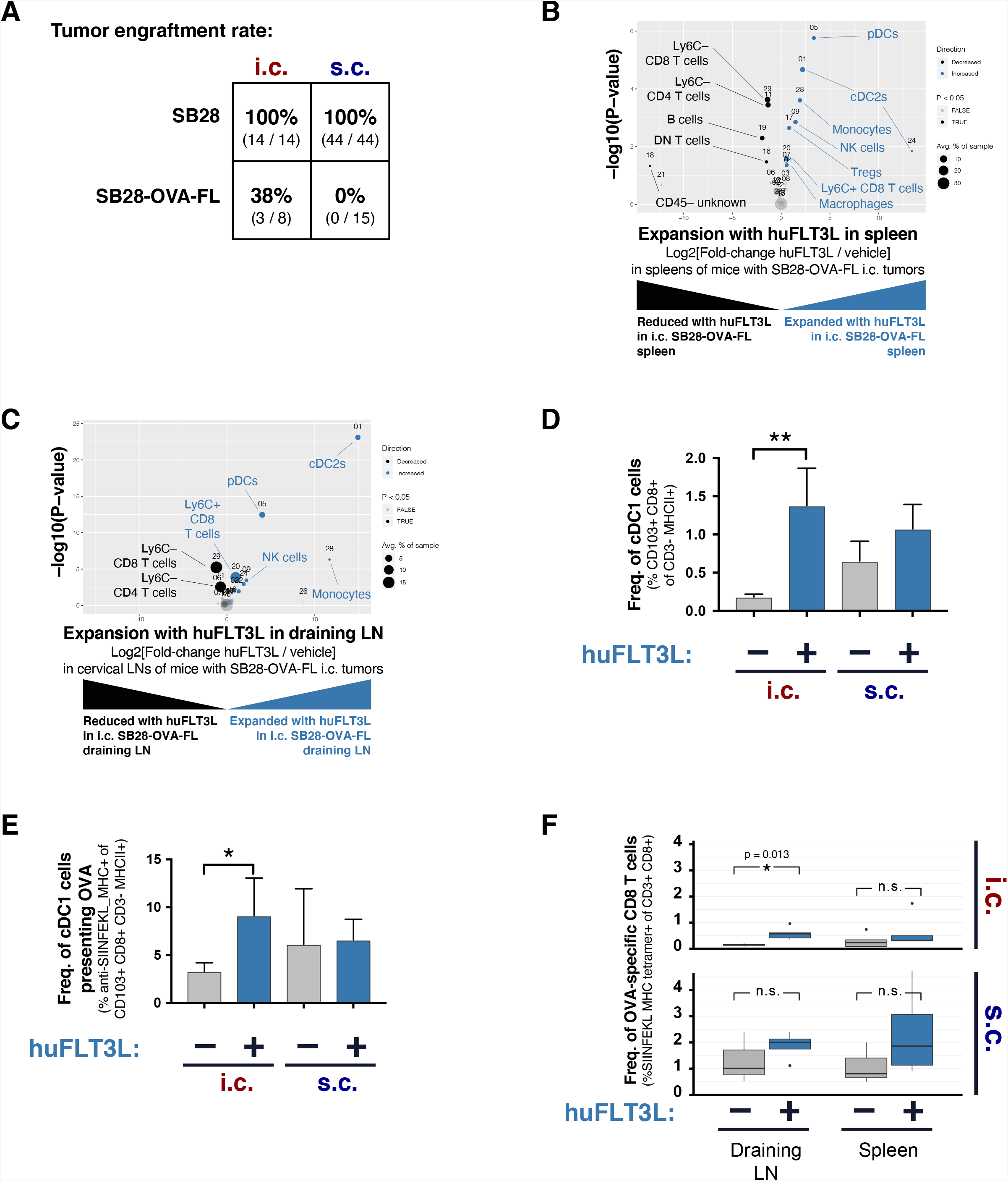
Treatment with FLT3L expands multiple dendritic cell subsets and improves antigen presentation in intracerebral SB28-OVA-FL tumors. **A**. Engraftment rates of SB28 (parental) and SB28-OVA-FL tumor lines in the intracerebral (i.c.) or subcutaneous (s.c.) sites. Data was aggregated from six independent experiments, details in **Suppl. Figure S4E.** **B**. Treatment with huFLT3L expands cDC2s, pDCs, and Tregs in the spleens of mice with SB28-OVA-FL tumors at Day 12 post-inoculation. Volcano plot comparing abundance of immune subpopulations in dissociated spleens from huFLT3L-treated (blue) and control-treated (black) intracerebral SB28-OVA-FL. Statistically significant clusters in volcano plots are highlighted in opaque color and indicated with a cell type label. **C**. Treatment with huFLT3L expands cDC2s and pDCs in the draining lymph node of mice with SB28-OVA-FL tumors at Day 12 post-inoculation. Volcano plot as in (B). **D**. Treatment with huFLT3L expands cDC1 cells in intracerebral but not subcutaneous SB28-OVA-FL tumors. Frequencies of cDC1 cells (CD103+ CD8+ as a frequency of MHCII+ CD3-events) were quantified by flow cytometry in tumor-draining lymph nodes of intracerebral and subcutaneous SB28-OVA-FL tumors +/− huFLT3L treatment. Tissues were harvested on Day 11 and 12 post-inoculation for s.c. and i.c. tumors, respectively. **E**. Treatment with huFLT3L improves presentation of a tumor-derived antigen. Frequency of cDC1 cells that were presenting SIINFEKL-peptide on MHC class I were quantified by flow cytometry with an anti-SIINFEKL-MHC-complex antibody. Same samples as in (D). **F**. Treatment with huFLT3L improves priming of tumor antigen-specific T cells. Frequency of OVA-specific CD8 T cells in tumor-draining lymph nodes or spleens of mice with intracerebral or subcutaneous SB28-OVA tumors +/− huFLT3L treatment, was quantified by flow cytometry with a SIINFEKL MHC-tetramer. Same samples as in (D).

### Systemic FLT3L expands cDC2s and pDCs in the SB28-OVA-FL intracerebral model

Results in the subcutaneous SB28-parental tumor model suggested that DCs play a critical role in establishing anti-tumor immunity, especially by priming CD4 T cells. Major DC subsets in tumors and tumor-draining lymph nodes can include: (1) plasmacytoid DCs (pDCs) that produce type I interferon in response to viral infection; (2) cDC1s that cross-present MHC-I epitopes to CD8 T cells; and (3) cDC2s that mediate priming of CD4 T cells via MHC-II^23^. Immune profiling showed that cDC2s were more abundant in the intracerebral GL261 TIME compared to intracerebral SB28 (**Figure 1D**), and cDC2s were expanded in SB28 subcutaneous tumors before and after CPI administration (**Figure 2D & 3C**), suggesting a role in CPI-mediated anti-tumor immunity. We hypothesized that increasing DC abundance at the intracerebral site might restore anti-tumor immunity. We therefore investigated if treatment with FLT3L, a key growth factor for DCs, would boost antigen presentation and T cell priming.

We inoculated mice either intracerebrally or subcutaneously with SB28-OVA-FL, and administered soluble half-life-extended (Fc-fused) human FLT3L (huFLT3L) daily for 10 days. Human FLT3L is cross-reactive with mouse FLT3 receptors and repeated injections can expand DCs in the periphery and in the brain^29^. We sacrificed the mice on Day 11-12 post-inoculation to perform mass cytometry profiling of splenocytes and tumor draining cervical lymph nodes. The systemic huFLT3L treatment caused expansion of cDC2s and pDCs (**Figure 4B,C** and **Suppl Figure S4A,B**) in spleens and draining lymph nodes. The expansion in draining lymph nodes was particularly dramatic, with pDCs increasing from 0.4% to 5.4%, and cDC2 expanding from 0% to 4.2%. The expansion of DC subsets was accompanied by upregulation of Ly6C+ on CD8 T cells (**Figure 4B,C** and **Suppl Figure S4A,B**). Induction of Ly6C on T cells is primarily driven by interferon-α signaling^30^, which may indicate that the expanded DCs were secreting type I interferons.

### Systemic FLT3L treatment improves antigen presentation in the SB28-OVA-FL model

Cross-presenting cDC1s (CD103+ CD8+) have a unique capability to prime CD8 T cells against MHC class I-restricted tumor antigens and have been shown to be essential for the effect of CPI in some tumor models^31,32^. We used flow cytometry to quantify CD103+ CD8+ double-positive cDC1s in the tumor-draining lymph nodes from intracerebral and subcutaneous SB28-OVA-FL tumors treated with either huFLT3L or vehicle (harvested Day 11-12). huFLT3L treatment robustly expanded the cross-presenting cDC1 population in draining lymph nodes from intracerebral SB28-OVA-FL, but had no significant effect on cDC1s in draining lymph nodes from subcutaneous tumors (**Figure 4D**).

We next measured the frequency of cDC1s that were cross-presenting the tumor-derived antigen SIINFEKL. Cells were stained with an antibody that recognizes SIINFEKL peptide docked in MHC-I and measured by flow cytometry. This assay confirmed that treatment with huFLT3L treatment significantly increased presentation of tumor antigen in intracerebral SB28-OVA-FL tumor-draining lymph nodes, matching the levels seen in subcutaneous tumor-draining lymph nodes (**Figure 4E**). Notably, huFLT3L treatment did not improve presentation of SIINFEKL on cDC1 in subcutaneous tumors (**Figure 4E**), just as it did not increase the frequency of cDC1 cells in those tumors (**Figure 4D**). To assess whether increased cross-presentation of tumor antigen led to increased priming of OVA-specific CD8 T cells, we stained cells with a SIINFEKL MHC-I tetramer. Indeed, huFLT3L treatment led to an expansion of OVA-specific CD8+ T cells in intracerebral SB28-OVA-FL tumor-draining lymph nodes (**Figure 4F**). Taken together, these results show that huFLT3L treatment induced expansion of cDC1, cDC2 and pDC populations in draining lymph nodes of intracerebral SB28-OVA-FL tumors, enhanced tumor antigen cross-presentation on cDC1s, and increased the frequency of tumor antigen-specific T cells.

### huFLT3L monotherapy extends survival in the SB28 intracerebral model

We then asked whether huFLT3L would improve survival in mice bearing intracerebral SB28 (parental) tumors which do not respond to dual CTLA-4/PD-1 blockade. All mice treated with an isotype control IgG died within 33 days of inoculation, while 5 of 22 mice (23%) receiving huFLT3L monotherapy lived more than 35 days (**Figure 5A**). Two of the huFLT3L-treated mice were effectively cured, surviving more than 80 days. We postulated that radiation (XRT) – a cornerstone of GBM clinical care – would promote priming by releasing tumor antigens for subsequently presentation by DCs. Therefore, we tested huFLT3L and fractionated high dose radiation therapy (XRT; 8 Gy x 3 doses), alone and in combination. The combination of huFLT3L + XRT improved survival relative to control-treated mice, but did not provide significant additional benefit over XRT or huFLT3L alone (**Figure 5B**). Paradoxically, combining huFLT3L treatment with dual CTLA-4/PD-1 blockade trended toward shorter survival (**Suppl. Figure 6A**). Altogether, these results support a small therapeutic benefit of huFLT3L treatment in SB28 intracerebral tumors. Given the multiple barriers to immune activation in SB28 intracerebral tumors, we explored more combination approaches in an effort to extend survival in a greater fraction of mice.

**Figure 5.**
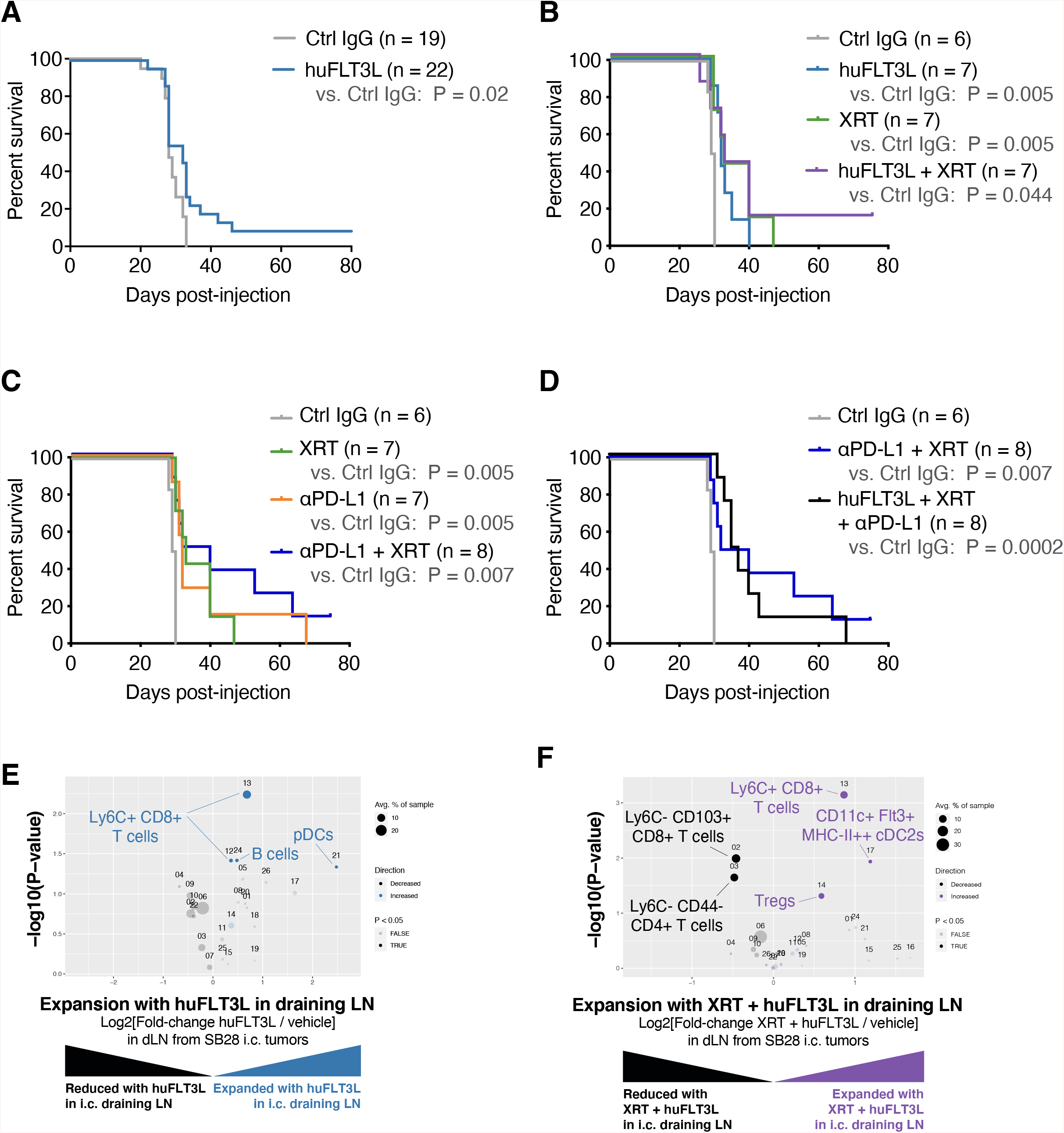
Improved antigen presentation combined with checkpoint blockade extends survival of mice with SB28 intracerebral tumors. **A**. Kaplan-Meier curves of intracerebral SB28 treated with IgG control (grey) or huFLT3L (blue). Log-rank test P-values are shown. Data from two experiments were aggregated. **B**. Kaplan-Meier curves of intracerebral SB28 treated with IgG control (grey), huFLT3L (blue), radiation (XRT, green) or huFLT3L and XRT (purple). Log-rank test P-values are shown. **C**. Kaplan-Meier curves of intracerebral SB28 treated with IgG control (same as B, grey), radiation (XRT, same ac B, green), anti-PD-L1 (yellow) or XRT and ant-PD-L1 (blue). Log-rank test P-values are shown. **D**. Kaplan-Meier curves of intracerebral SB28 treated with double (anti-PD-L1 and XRT, same as C, blue) or triple-combination immunotherapy (huFLT3L and XRT and anti-PD-L1, black). Log-rank test P-values are shown. **E**. Volcano plot comparing abundance of immune subpopulations in draining lymph nodes from intracerebral SB28 treated with huFLT3L (blue) versus control (black) using CyTOF. Statistically significant clusters in volcano plots are highlighted in opaque color and indicated with a cell type label. **F**. Volcano plot as in (E) comparing radiation (XRT) + huFLT3L (purple) versus control (black).

### PD-L1 blockade extends survival in the SB28 intracerebral model

Our immune profiling of patient samples and mouse models indicated an overabundance of PD-L1+ TAMs in GBM. While dual CTLA-4/PD-1 blockade was ineffective in SB28 intracerebral tumors (**Figure 2B**), PD-L1 blockade slightly extended median survival relative to isotype control IgG-treated tumors (32 vs. 29.5 days, P = 0.005) (**Figure 5C**). This effect was further enhanced by addition of XRT (36 vs. 29.5 days, P = 0.007). Addition of huFLT3L (i.e. triple combination therapy) did not improve survival relative to PD-L1 + XRT (**Figure 5D**). We next sought to enhance the maturation or activation of DCs and tested a CD40 agonistic antibody or polyIC:LC (TLR3 agonist) in combination with huFLT3L. Addition of CD40L to huFLT3L treatment trended toward providing additional benefit, but did not significantly improve survival relative to huFLT3L alone (**Suppl Figure 6D**). PolyIC:LC did not provide any additional benefit relative to huFLT3L alone (**Suppl Figure 6D**).

### huFLT3L expands DCs and promotes T cell maturation in the SB28 intracerebral model

Mass cytometry of tumor-draining lymph nodes after huFLT3L monotherapy showed expansion of pDCs as well as upregulation of Ly6C+ on CD8+ T cells, but there was no change in total CD8+ T cell numbers (**Figure 5E and Suppl. Figure S5A**). While expansion of the pDCs is likely a direct consequence of huFLT3L treatment, the upregulation of Ly6C+ T cells may be a byproduct of interferon-α production by the pDCs^30^. The dual therapy of XRT+hFLT3L had similar effects, but produced fewer pDCs and more cDC2s. This combination therapy also caused a 1.5-fold expansion of Tregs coupled with a 1.5-fold reduction of naive (CD44-) CD4 T cells (**Figure 5F**). Similar increases in Tregs were observed when huFLT3L was combined with dual CTLA-4/PD-1 blockade (**Suppl Figure 6B**). Taken together, these results suggest that FLT3L combination immunotherapies can promote a pro-inflammatory microenvironment in draining lymph nodes, at the cost of a compensatory expansion of Tregs that may ultimately contribute to immune evasion.

As for the immune compartment within SB28 intracerebral tumors themselves, XRT monotherapy reduced cDC2s and expanded a PD-L1+ CD206+ TAM population, both of which would be undesirable changes for anti-tumor immunity (**Suppl Figure S5B & S5D**). Combining huFLT3L treatment with XRT reversed these effects, potentially creating a tumor microenvironment more supportive of T-cell priming (**Suppl Figure S5C**). Neither XRT monotherapy, huFLT3L monotherapy, nor the combination altered the abundance of intratumoral T cells at endpoint (data not shown).

We then investigated whether these treatments cause a change in surface marker profiles on intratumoral T cells. Normal human brain contains rare CD69-high CD8+ T cells, a type of tissue-resident memory cells (T_RM_) ^33^. We also observed that the CD8+ T cells in untreated tumors were CD69-high CCR7-low PD-1-high (**Suppl. Figure S6E**). Both XRT and huFLT3L monotherapies, as well as the combination, promoted a shift in the phenotype of intratumoral CD8+ T cells characterized by downregulation of PD-1 and CD69 and upregulation of CCR7 (**Suppl. Figure S6E**). This change in T cell surface marker expression, especially the upregulation of CCR7, may suggest differentiation of T_RM_ cells toward a central memory T cell fate.

In summary, FLT3L treatment, blockade of PD-L1 and radiation all improved survival in SB28 intracerebral tumors. There was a benefit of combining PD-L1 blockade with radiation, but triple combinations did not provide additional survival benefit. Each treatment or combination impacted the immune microenvironment in the tumor and draining lymph node in slightly different ways, primarily affecting T cell and dendritic cell subsets. Our data suggest that that DCs and PD-L1+ TAMs are malleable cell populations in human and mouse gliomas that can be modulated therapeutically using huFLT3L, anti-PD-L1 and radiation.

## DISCUSSION

Our study shows that intracerebral GBM tumors fail to elicit anti-tumor immunity due to defects early in the cancer-immunity cycle^34^, specifically, poor priming of antigen-specific T-cells at the tumor-draining lymph node. Additionally, there may be defects later in the cycle mediated by PD-L1+ cells in the TIME, as suggested by the efficacy of PD-L1 blockade.

Although PD-1 and PD-L1 checkpoint blockade therapies disrupt the same signaling axis, our results indicate that PD-L1 blockade is more effective than dual CTLA-4/PD-1 blockade in the SB28 model. This is in line with a recent report of a modest positive effect in interim data from a phase II clinical trial combining standard XRT and PD-L1 blockade in unmethylated GBM^35^. SB28 appears to be a faithful model of PD-L1 expression and myeloid infiltration in human GBM. In a previous study of the GBM microenvironment by flow cytometry, myeloid cells were proposed as the dominant source of PD-L1^36^. Consistent with these results, our mass cytometry profiling showed that PD-L1-expressing myeloid cells were abundant in human GBM as well as the SB28 model. It has been shown that SB28 glioma cells do not express PD-L1 under basal conditions, but PD-L1 expression can be induced *in vitro* by IFNg^22^. Why anti-PD-L1 was more effective than dual CTLA-4/PD-1 blockade is unknown, but there are several possible mechanisms that distinguish these therapies. Anti-PD-L1 is able to disrupt the *cis* interaction between PD-L1 and B7-1 (CD80) on DCs, allowing CD80 to activate T cells via CD28^37^. Anti-PD-L1 is also able to act directly on tumor cells, driving cytokine production and *in vivo* phagocytic activity of glioma TAMs in some contexts^38^. Finally, dual CTLA-4/PD-1 blockade can induce apoptosis of tumor-specific T cells in preclinical models with low tumor burden^39^.

Several lines of evidence suggest that defective antigen presentation is of central importance in explaining the non-responsiveness of GBM to CPI. Our results in the SB28 model agree with recent reports that brain tumors are poorly infiltrated by DCs^9,10^. We observed dramatic differences in antigen presentation and responsiveness to immunotherapy when SB28 tumors were grown in the flank as opposed to the brain. Expression of multiple strong MHC class I and II-restricted antigens in SB28 tumor cells (SB28-OVA-FL) completely prevented engraftment in the flank, but only reduced engraftment in the brain by about two-thirds. SB28 flank tumors showed a significant influx of cDC2s into the tumors and cDC1s into the tumor draining lymph nodes, as compared to SB28 intracerebral tumors. This influx was further increased by dual CTLA-4/PD-1 blockade. Further supporting a central role for DCs, the dual CTLA-4/PD-1 blockade-mediated inhibition of SB28 flank tumors was dependent on CD40 signaling, which is important for licensing of DCs. Treatment with huFLT3L increased the frequency of OVA-presenting cDC1s in cervical lymph nodes and modestly improved survival, thus highlighting the fundamental defects in antigen presentation in the brain, and providing a rationale for FLT3L-based strategies to overcome this challenge. This is also in line with a recent study in which dual CTLA-4/PD-1 blockade promoted rejection of melanoma brain metastases only when an extracranial melanoma tumor was present as well^40^. Another recent report showed that intracranial delivery of VEGF-C can boost antigen trafficking to deep cervical lymph nodes and drive rejection of GL261 tumors^41^. This is a promising new way to modulate antigen trafficking and an exciting candidate for follow-up in the SB28 model in combination with the approaches tested here. Overall, boosting antigen presentation will be an important component in an effective brain immunotherapy regimen.

The dual CTLA-4/PD-1-mediated rejection of SB28 flank tumors was dependent on CD4 T cells and NK cells, while CD8 T cells were less important, suggesting an unconventional effector mechanism of anti-tumor immunity. In our hands, SB28 tumor cells did not express MHC-II, consistent with previous reports^22^, so it is unlikely that effector CD4 T cells are directly mediating tumor rejection in this model. Based on our depletion data, the critical effector cells in CPI-treated SB28 subcutaneous tumors are likely NK cells that are potentiated by antigen-specific CD4 T cells. NK cells and CD4 T cells were also expanded over time in untreated subcutaneous SB28 tumors, indicating they are involved in tumor surveillance. CD4 T cells were recently implicated as critical drivers of anti-tumor immunity in a mouse model of breast cancer^42^, and CD4 T cells – but not CD8 T cells or NK cells – were critical for eliciting anti-tumor immunity in a rat model of glioma^43^. The essentiality of CD4 and NK cells in the SB28 model is particularly interesting in light of three other findings: (1) priming in the periphery elicited systemic immunity, (2) the efficacy of dual CTLA-4/PD-1 blockade in the flank required CD40-mediated licensing of DCs, and (3) cDC2 cells were abundant even in untreated SB28 flank tumors. Integrating these pieces of evidence, we propose a model of the systemic anti-tumor immunity that we observed in SB28 flank tumors treated with dual CTLA-4/PD-1 blockade: Tumor antigen is processed and presented by cDC2s early in the life of the tumor, but it is not sufficient to activate CD4 T cells due to negative signals from PD-1 and CTLA-4. The addition of dual CTLA-4/PD-1 checkpoint blockade expands cDC2s and lowers the threshold of activation, promoting CD4 T cell expansion and release of pro-inflammatory cytokines (e.g. IL-2, IFN gamma, IL-12). These cytokines, in turn, promote expansion and activation of NK cells^44^, which act as the main effector cell population to mediate SB28 flank tumor rejection. We postulate that memory CD4 T cells are also key mediators of systemic immunity in this model, protecting mice from intracerebral rechallenge by secreting pro-inflammatory cytokines and activating NK cells at the new tumor site.

In SB28 intracerebral tumors, FLT3L treatment led to a modest but significant increase in overall survival, including sustained remissions of over 60 days in a subset of mice. However, combining FLT3L with other agents or radiation did not lead to synergistic or additive effects. To harness the potential of FLT3L treatment we need to understand what distinguishes the few cured mice from the non-responders and why combination of dual CTLA-4/PD-1 blockade with FLT3L appeared to negate the benefits of FLT3L treatment. Evidence from our mass cytometry profiling points to regulatory T cells as a key player in this system. We observed that FLT3L monotherapy caused expansion of pDC and Tregs in tumor-draining lymph nodes, which was further enhanced when FLT3L was combined with XRT or dual CTLA-4/PD-1 blockade. The mechanism underlying this expansion of Tregs may be similar to that previously described in human breast and ovarian cancer^45,46^ and the B16 mouse model of melanoma^31^. In those studies, tolerogenic pDCs engaged ICOS on CD4 T cells, thereby promoting differentiation to Tregs, and causing immunosuppression in tumors. Similarly, Tregs and cDC2 numbers were shown to be positively correlated in the B16 melanoma model, and Treg depletion could restore CD4 T cell-mediated immune rejection^47^. Thus, our data suggest that expansion of DCs with FLT3L in the SB28 intracerebral model can initiate homeostatic feedback mechanisms that blunt the efficacy of FLT3L as a monotherapy or in combination with CPI.

Based on our results here, FLT3L shows potential as a therapeutic strategy to improve antigen presentation in GBM, but combination strategies will likely be necessary to achieve a clinical benefit. Human clinical trials using FLT3L in mesothelioma and some carcinomas have reported limited effect as a single agent^48,49^. However, pre-clinical studies in several mouse tumor models suggest that FLT3L can be effective, particularly when used in combination with other agents. In a mouse model of pancreatic adenocarcinoma, a combination of FLT3L, XRT and CD40 agonist was used to skew the DC phenotype away from a Th2 and Th17 response toward a Th1 response^50^. In a mouse model of BRAF-mutant melanoma, CD103+ cDC1s were essential for an effective anti-PD-L1 response and could be expanded by FLT3L. However, FLT3L treatment also led to proliferation of immature progenitor DCs with the capacity to expand Tregs. Combining FLT3L with an agent to promote maturation of DCs (poly I:C) was essential to promote an effective anti-PD-L1 mediated immune response^31^. These examples were all done in extracranial tumor models, but in the intracerebral SB28 model, the addition of polyIC:LC or CD40 agonist did not substantially improve FLT3L therapy.

Overall, this work illustrates how tumor location, antigenicity, and immune microenvironment can influence the efficacy of immunotherapies. It underscores how these factors must work in harmony to tip the scales toward effective antigen presentation and anti-tumor immunity. Our data demonstrate that the main hindrance for effective immunotherapy of glioma is poor antigen presentation, rather than intrinsic immunosuppressive properties of the glioma cells themselves. Accurate syngeneic tumor models and deep immune profiling should be leveraged to further elucidate the immunosuppressive mechanisms that are unique to brain tumors, and to guide the rational combination of immunotherapies to control these challenging tumors.

## DECLARATIONS

### Availability of data and material

Mass cytometry data will be made publicly available on FlowRepository upon publication. Analysis scripts will be made publicly available on GitHub upon publication.

### Authors’ contributions

Experimental design, generation, and analysis of data: E.F.S, E.D.L, L.K.M., N.M.N., P.C., A.S., V.G., P.R.W., F.J.S., M.H Manuscript figure assembly: E.F.S, E.D.L, Reagents: P.B. Patient material and data: E.D.L., E.L., W.T., C.R., J.S., M.S., L.F., S.A.S., L.D.W., C.E.B. Supervision: L.F., C.E.B., H.O, W.A.W, M.H. Manuscript writing: E.F.S, H.O, W.A.W, M.H.

## Acknowledgements

We thank: Anny Shai and Joanna Phillips at the UCSF Brain Tumor Research Center for banking of glioma samples. The staff at the UCSF Helen Diller Family Comprehensive Cancer Center Preclinical Therapeutics Core for assistance with mouse studies. Stanley Tamaki, Michael Lee, and Claudia Bispo at the UCSF Single Cell Analysis Center for assistance with mass cytometry studies.

