## Supplemental Methods for "Deep immune profiling reveals targetable mechanisms of immune evasion in checkpoint blockade-refractory glioblastoma"

### Patient Samples

All procedures performed in studies involving human participants were in accordance with the ethical standards of the institutional and/or national research committee and with the 1964 Helsinki declaration and its later amendments or comparable ethical standards.

Patients provided written and informed consent to tissue collection under UCSF institutional review board (IRB) protocol number 10-01318 for brain tumors or 13-12246 for kidney and sarcoma tumors. One additional brain tumor (COH001) was collected with informed consent under City of Hope IRB Protocol Number 07074.

Eighteen (18) deidentified human glioblastoma specimens were obtained from the University of California, San Francisco (UCSF) Brain Tumor Research Center and one from City of Hope (COH) (**Suppl Table S1**). Fourteen (14) human renal cell carcinoma (RCC) specimens (including 1 metastatic tumor biopsy and 1 normal adjacent tissue biopsy) and five (5) human sarcoma samples (including 1 normal adjacent tissue biopsy) were collected from patients at UCSF (**Suppl Table S1**). Dissociated, viably cryopreserved samples were thawed, stained with cisplatin (Enzo) and fixed with paraformaldehyde as described below.

### Cell lines, reagents

Cell lines (SB28-Parental and GL261) were grown in DMEM with 10% FBS. All cells were tested and authenticated. Mycoplasma status was checked monthly using PlasmoTest Mycoplasma Detection Kit (InvivoGen). Cells were passaged every 3-5 days with TrypLE Express Enzyme (Life Technologies) when confluent and re-plated in T75 flasks at 1e5-5e5 cells in 15 mL media.

### In vivo studies

Animal experiments were performed in accordance with national guidelines and regulations, and UCSF IACUC Protocol Numbers AN169783-01C or AN152965-02A. For all intracranial and flank tumor studies, 7-week-old C57BL/6 mice were acquired from Jackson Laboratory (stock number 000664) and acclimated for 7 days prior to injection date such that they were 8 weeks old on the day of tumor injection (day 0). Tumor-bearing mice were monitored for tumor growth 1-2 times weekly via bioluminescence imaging using an IVIS Spectrum in vivo imaging system (PerkinElmer). Imaged mice received intraperitoneal injection of potassium-salt D-luciferin (GoldBio) diluted in sterile saline at 20 mg/mL, for a dose of 64 mg/kg, 20 minutes prior to measurement of radiance. Tumor volume of subcutaneous flank tumor-bearing mice was measured 1-2 times weekly via caliper measurement, and volume was approximated by the modified ellipsoid formula (length x width^2)/2. Endpoint criteria by which mice were euthanized were defined per our approved IACUC protocol as exhibiting > 15% weight loss from baseline weight taken on the day of injection, and/or exhibiting signs of pain (i.e. hunch, grimace), ulceration of the skin overlying subcutaneous tumors without improvement over a period of 7 days, tumor width exceeding 2 centimeters, abnormal neurological signs (such as seizures), or decreased motility.

### Mouse intracranial injections

8-week-old C57BL/6 mice were used for intracranial studies. Cell lines (GL261, SB28-Parental, SB28-OVA-FL) were trypsinized with TrypLE (Life Technologies) and washed with DMEM and sterile saline before being resuspended in DMEM for inoculation. Mice were anaesthetized with isoflurane and injected orthotopically with SB28-Parental cells in 2 uL DMEM containing 50,000 or 1,600 cells as specified in **Suppl Table S5**. Using a stereotactic frame, a burr hole was formed on the skull via 0.7 mm drill bit 1.5mm laterally to the right and 1.5mm rostrally from the bregma, and a non-coring needle (Hamilton 7804-04, 26s gauge, Small Hub RN Needle, Point Style: 4, Needle Length: 2 inches, Angle: 12) was used to inject the cells at a depth of 3mm into the brain from the burr hole. The burr hole was covered with bone wax, and the skin incision was sutured. Mice were given an intraperitoneal injection of buprenex (0.05 mg/kg) once directly after surgery and again 4-6 hours post-surgery, an intraperitoneal injection of meloxicam injection before surgery and 12-24 hours post-surgery, and monitored for signs of pain. Only mice with no signs of pain after 24 hours post-surgery were used for study.

### Mouse subcutaneous flank injections

8-week-old C57BL/6 mice were injected subcutaneously at the right-sided flank region using a sterile 0.5cc insulin syringe with 5e5 cells (SB28-Parental or SB28-OVA-FL) suspended in 100 uL of DMEM or 1:1 DMEM:Matrigel (Corning) according to **Suppl Table S5**.

### Treatment regimens and dosing

Therapeutic antibodies, depletion antibodies, recombinant cytokines, and poly-IC:LC were diluted with sterile saline and injected according to the doses and treatment schedules in **Suppl Table S6**. Poly-IC:LC (Hiltonol) was kindly provided by Dr. Andres M. Salazar (Oncovir Inc, Washington, DC).

A small animal radiation research platform (SARRP, Xstrahl Inc.) was used to administer radiation (XRT) in intracranial tumor-bearing mice. Mice were placed under isoflurane anesthesia via nose-cone adapter on the SARRP platform. The burr hole was visualized via CT scan, and the isocenter of XRT was set to 3 mm deep to the burr hole on the skull, targeting the site at which the tumor was injected. Radiation was administered via a 3mm square collimator in a 90° arc (incidence angle of -45° to 45° laterally about the dorsoventral axis) according to the doses and treatment schedules in **Suppl Table S6**.

### Dissociating mouse and human samples for CyTOF

For intracranial tumors, mice were first perfused via intracardiac injection of 10 mL sterile PBS using a perfusion pump before harvesting the whole brain and dissection of the whole tumor mass at 4°C. Tumors were immediately minced in 10 cm dishes with sterile scalpels. Tumor fragments were washed with PBS, transferred to capped tubes, and enzymatically digested with a cocktail of collagenase IV (Worthington, 3.2 mg/mL), deoxyribonuclease I (Worthington, 1 mg/mL), and soybean trypsin inhibitor (Worthington, 2 mg/mL) diluted in sterile PBS. 600 uL of this cocktail was used per 100 mg of tumor. Tumor fragments were incubated in a temperature-controlled rotating hybridization oven at 37°C, with additional mechanical disruption by trituration with a P1000 pipet tip 30 times every 15 minutes. After 45 minutes of digestion, cells were filtered through a 70 µm cell strainer and washed with sterile PBS. Residual erythrocytes were lysed with ACK Lysing Buffer (ThermoFisher) for 5 minutes before quenching with a 9-fold excess of PBS. Cells were then washed with PBS, counted, and stained with cisplatin (Enzo, 5 µM) for 5 minutes ta room temperature. Cisplatin staining was quenched by adding 5 mL of AutoMACS Running Buffer (Miltenyi) AutoMACS Running Buffer (Miltenyi). Cells were washed, resuspended in AutoMACS Running Buffer, and fixed by adding methanol-free paraformaldehyde (PFA) (Electron Microscopy Sciences, 1.6%) for 10 minutes. Cells were washed, resuspended in AutoMACS Running Buffer plus 10% DMSO at a density of 2e6-1e7 cells/mL, then frozen at -80°C.

Mouse spleens and lymph nodes were harvested from euthanized mice and immediately placed on ice. Spleens were gently mashed through a 70 µm cell strainer (Fisher Scientific) while lymph nodes were gently mashed through a 35µm cell strainer (Fisher Scientific) in PBS. Erythrocytes were lysed and cells were stained with cisplatin and fixed with PFA prior to freezing, as above.

### CyTOF antibody conjugation and validation

Metal-tagged antibodies were purchased pre-conjugated (Fluidigm Corporation, South San Francisco, CA) or conjugated in-house using the MAXPAR™ X8 chelating polymer kit (Fluidigm Corporation, South San Francisco, CA). Some metal isotopes were obtained from Trace Sciences International (Richmond Hill, Ontario, Canada) and were prepared by the UCSF Single Cell Analysis Center (UCSF SCAC). The concentration of final conjugated antibody stocks were measured by 280 nm absorbance on a NanoDrop and diluted to a maximum concentration of 0.8 mg/mL for storage at 4°C. Metal-tagged antibody stocks were validated and titrated using positive-control and negative-control cells on the Fluidigm Helios™ mass cytometer at UCSF SCAC. Antibody panels are detailed in **Suppl Tables S3, S4, and S5**.

### Staining of mouse and human samples with CyTOF antibodies

Mouse tumor, spleen, and lymph node samples were thawed, counted, and stained 24-72 hours prior to data acquisition on the Helios mass cytometer. Samples were barcoded with Fluidigm Cell-ID 20-Plex Pd Barcoding Kit according to the manufacturer's protocol. Barcoded samples were pooled then washed by centrifuging at 600 x *g* 5 mins and resuspending in AutoMACS Running Buffer. For mouse samples, Fc blocking was performed with anti-mouse CD16/CD32 antibody (clone 2.4G2) (UCSF SCAC) at 1 ug per 1e6 cells for 10 mins at room temp. For human samples, Fc blocking was performed with TruStain FcX (BioLegend) at 8 µL per 100 µL of cell suspension for 5 mins at room temp. Mouse samples were stained with FITC-conjugated anti-CCR2 and biotin-conjugated anti-CCR4 for 30 mins, then washed twice with AutoMACS Running Buffer. Cells were then stained with metal-tagged antibodies against surface-expressed targets for 30 mins at room temp, then washed twice with AutoMACS Running Buffer. Surface-stained cells were resuspended in 3 mL of Perm-S Buffer (Fluidigm) and incubated for 20 min at room temp (human samples) or 30 min at 4°C (mouse samples) for permeabilization. Permeabilized cells were centrifuged and resuspended in Perm-S Buffer with a cocktail of antibodies against intracellular markers, then incubated 30 mins at room temp. Cells were washed twice with Perm-S Buffer and twice with AutoMACS Running Buffer. Mouse samples were intercalated and post-fixed at 4°C up to 7 days in PBS with 7% Perm-S buffer (v/v), 0.1µM Ir intercalator (Fluidigm), and 4% PFA. Human samples were intercalated and post-fixed at 4°C up to 7 days in PBS with 80% Perm-S buffer (v/v), 0.25µM Ir intercalator, and 3% PFA. Cells were centrifuged and resuspended in 18MΩ water with EQ™ Four Element Calibration Beads (Fluidigm) prior to running on the Fluidigm Helios™ mass cytometer at UCSF SCAC.

### CyTOF data acquisition and analysis

Data was acquired on a Fluidigm Helios™ mass cytometer. Data was preprocessed in six steps:

(1) Raw linear data was asinh transformed with a cofactor of 5, as previously described (Bendall et al., *Science* 2011).

(2) Measured channels were normalized over a sliding time window, using a reference signal from metal-embedded beads. The same normalization algorithm was used across datasets, but it was implemented in two different software packages:

   a. All Mouse panels and Human Myeloid panel FCS files were normalized using a Matlab-based normalization software (Finck et al *Cytometry A* 2013).

   b. Human T-cell panel FCS files were normalized using the R-based debarcoder in the CATALYST suite (Nowicka et al *F1000Research* 2017).

(3) Viable cells were debarcoded to identify singlet cells with valid palladium-based barcodes. The same debarcoding algorithm was used across datasets, but it was implemented in two different software packages:

a. All Mouse panels and Human Myeloid panel FCS files were debarcoded using the Matlab-based debarcoder (Zunder et al *Nat Protocol* 2015) and a global separation threshold of 0.3.

b. Human T-cell panel FCS files were debarcoded using the R-based debarcoder in the CATALYST suite (Nowicka et al *F1000Research* 2017). Global cutoff of 0.2.

(4) Channel name metadata in the FCS file was corrected and harmonized, when necessary, using the cytutils package for R (github.com/ismms-himc/cytutils).

(5) Signal from all palladium barcode channels was summed and written as a new synthetic channel called "Pd_sum". This channel served as an approximation of total cell protein content, and provided a useful counterstain for manually gating dead cells in the following step.

(6) Data was manually gated in Cytobank to remove dead cells labeled with cisplatin (Fienberg et al *Cytometry A* 2012).

FCS files containing debarcoded viable singlet cells were used as input to "PhenoSOM", a clustering and visualization script written in the R programming language for this project. PhenoSOM links together published R packages to perform three major steps:

(1) For each FCS file (e.g. patient sample), high-resolution clusters were identified by clustering all cells into 900 SOM nodes (30 x 30 grid) using the FlowSOM package for R (van Gassen et al *Cytometry A* 2015). This reduced the input to the PhenoGraph algorithm from millions of cells to hundreds of clusters, dramatically speeding it up. Channels used for clustering are indicated in **Suppl Tables S3, S4 and S5.**

(2) Metaclusters were identified by aggregating the SOM nodes (900 per file) from all files in the dataset, and using this as input to the PhenoGraph algorithm (Levine et al *Cell* 2015). A user-defined *k* parameter (number of nearest neighbors) between 10 and 30 was chosen by manual inspection of cluster size and stability. The R implementation of the PhenoGraph algorithm, part of the Cytofkit R package (Chen et al *PloS Computational Biology* 2016), was used.

(3) To identify lymphocyte subpopulations (e.g. tumor-immune microenvironment; TIME) while excluding non-immune cell types, the PhenoGraph metaclusters with above-background median CD45 expression were selected for input into a second round of clustering. The threshold for CD45 expression was between 0.4 and 1.0 (asinh-transformed values), depending on the dataset. For each original FCS file (e.g. patient sample), the CD45-positive cells were used as input to a second round of FlowSOM clustering (10 x 10 grid).

(4) TIME metaclusters were identified by aggregating the SOM nodes from the second round of clustering (100 per file) from all files in the dataset, and using this as input to the PhenoGraph algorithm, as described above.

(5) Comparisons between different experimental conditions were made using the edgeR algorithm for differential expression (Robinson et al *Bioinformatics* 2010) to assess changes in TIME metacluster abundance between groups of samples. The output of the edgeR algorithm was expressed as a volcano plot (fold-change vs. P-value).

The PhenoSOM pipeline (FlowSOM nodes → PhenoGraph metaclusters → diffcyt differential abundance analysis) is, to our knowledge, novel. However, bears similarities to the pipeline proposed by Nowicka et al (Nowicka et al *F1000Research* 2017). The main difference is that Nowicka et al used the ConsensusClusterPlus algorithm to identify metaclusters, instead of PhenoGraph. Given their similarities, we reasoned that these approaches should give qualitatively similar results if run on the same dataset, although there would be likely differences in cluster boundaries and levels of significance. We therefore repeated the differential abundance analysis from Suppl. Figure S5B using the Nowicka et al pipeline. This pipeline identified 1 cluster that was significantly enriched in XRT-treated tumors and matched the phenotype of the cluster identified by the PhenoSOM pipeline (**Suppl. Figure S5D**).

Immune populations in human and mouse CyTOF datasets were interpreted by manually inspecting the heatmaps of marker median expression in PhenoSOM metaclusters. PhenoSOM metaclusters were annotated with immune cell phenotypes by manual inspection of marker medians.

### Flow cytometry

Fluorescent flow cytometry data was acquired on a Becton-Dickinson LSR-II. Splenocytes or lymph nodes were dissociated in a 70 µm mesh and stained with DyLight 800 NHS Ester (ThermoFisher, 300 ng/mL, 15 min at 37°C) to permanently label dead cells. Cells were washed, fixed, and frozen as above. After thawing, Fc receptors were blocked with TruStain FcX anti-mouse CD16/32 FcX (BioLegend, 20 µg/mL, 10 minutes at 20°C). Fluorescent-conjugated antibodies or tetramers were added at the concentrations listed in **Suppl Table S7.**

### Hierarchical clustering of gene expression data

All RNA sequence reads were mapped against the mouse genome assembly mm9 using STAR v2.5.3a. The alignment was performed with a two-pass approach where splice junctions discovered during a first alignment guides the forming of a final second alignment (Veeneman et al *Bioinformatics* 2016). Unmapped reads were realigned using Bowtie2 v2.3.4.3 before duplicate reads were removed with Piccard v2.10.3. The expression profile was quantified with Subread v1.5.2 and TPM normalized using R v3.6.0.

Two gene expression datasets were downloaded to illustrate tumor type and subtype context. Brain tumors and normal brain from Gene Expression Omnibus using GEO accession GSE50161 (https://www.ncbi.nlm.nih.gov/geo/) and the TCGA-GBM micro array cohort from Genomic Data Commons (https://portal.gdc.cancer.gov/legacy-archive/). The reference data were downloaded as raw CEL files and preprocessed and RMA normalized using the R package oligo and hgu133plus2.db. Hierarchical clustering were performed using the Metagene code for cross-platform, cross-species characterization of global transcriptional states (Tamayo et al *PNAS* 2007).

### Transfection of SB28 with full-length OVA and OVA-II

To establish an SB28 cell line expressing the full-length OVA peptide (SB28-OVA-FL), pAC-Neo-OVA from Addgene (#22533) was used. Cells were passaged multiple times under selective antibiotic pressure with 500ug/ml G418 in DMEM with 10% FBS, and expression of peptide was confirmed via western blot with antibody (ab17293) from Abcam.

### Statistical analysis

Optical flow cytometry data was manually gated in Cytobank. Statistical analyses of manually gated flow cytometry population frequencies and *in vivo* data were performed in GraphPad Prism.

### Data accessibility

Processed mass cytometry data with PhenoGraph cluster assignments will be deposited online at FlowRepository.org upon publication.
