## Supplemental Figures S1-S6 for "Deep immune profiling reveals targetable mechanisms of immune evasion in checkpoint blockade-refractory glioblastoma"

### Figure S1

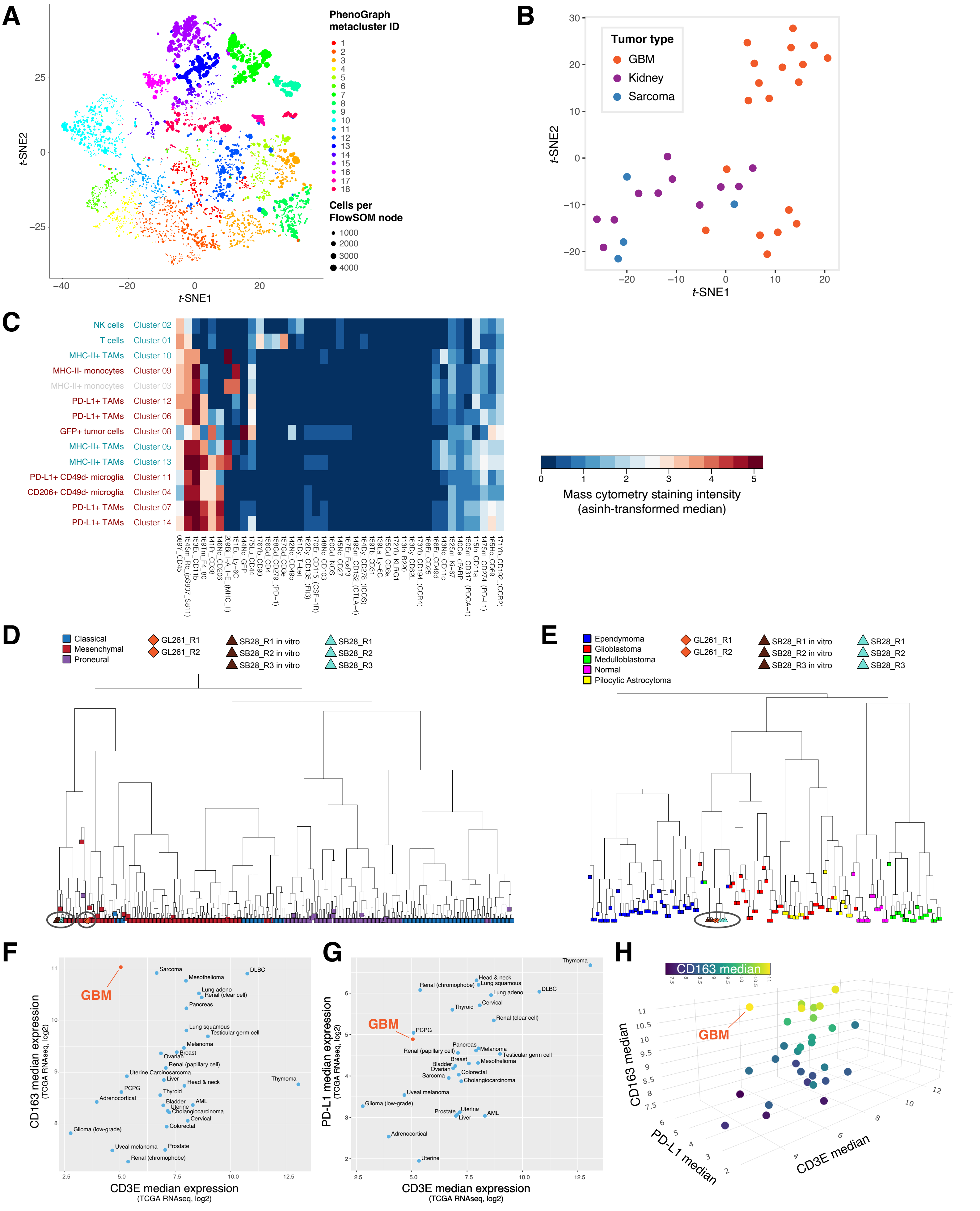

A
PhenoGraph metacluster ID
Cells per FlowSOM node
1000
2000
3000
4000
t-SNE1
t-SNE2
B
Tumor type
GBM
Kidney
Sarcoma
t-SNE1
t-SNE2
C
NK cells Cluster 02
T cells Cluster 01
MHC-II+ TAMs Cluster 10
MHC-II- monocytes Cluster 09
MHC-II+ monocytes Cluster 03
PD-L1+ TAMs Cluster 12
PD-L1+ TAMs Cluster 06
GFP+ tumor cells Cluster 08
MHC-II+ TAMs Cluster 05
MHC-II+ TAMs Cluster 13
PD-L1+ CD49d- microglia Cluster 11
CD206+ CD49d- microglia Cluster 04
PD-L1+ TAMs Cluster 07
PD-L1+ TAMs Cluster 14
Mass cytometry staining intensity (asinh-transformed median)
D
Classical
Mesenchymal
Proneural
GL261_R1
GL261_R2
SB28_R1 in vitro
SB28_R2 in vitro
SB28_R3 in vitro
SB28_R1
SB28_R2
SB28_R3
E
Ependymoma
Glioblastoma
Medulloblastoma
Normal
Pilocytic Astrocytoma
GL261_R1
GL261_R2
SB28_R1 in vitro
SB28_R2 in vitro
SB28_R3 in vitro
SB28_R1
SB28_R2
SB28_R3
F
GBM
CD163 median expression (TCGA RNAseq, log2)
CD3E median expression (TCGA RNAseq, log2)
G
GBM
PL-L1 median expression (TCGA RNAseq, log2)
CD3E median expression (TCGA RNAseq, log2)
H
CD163 median
GBM
CD163 median
PD-L1 median
CD3E median

### Supplementary Figure Legends

#### Figure S1.

A) Same as Figure 1A except Myeloid CyTOF panel

B) Same as Figure 1A except Myeloid CyTOF panel

C) Heatmap of cluster medians corresponding to volcano plot in Figure 1D

D) Gene expression profiles of GL261 orthotopic tumors (n=2), SB28 orthotopic tumors (n=3), or SB28 cells grown in vitro (n=3) were compared with primary human gliomas from Verhaak et al 2017 by hierarchical clustering.

E) Gene expression profiles of GL261 orthotopic tumors (n=2), SB28 orthotopic tumors (n=3), or SB28 cells grown in vitro (n=3) were compared with primary human brain tumors and normal brain tissue (n = 13) by hierarchical clustering.

F) TCGA analysis of human GBM shows an immunosuppressive TME characterized by a higher ratio of M2 TAMs to T cells (CD163:CD3), as compared to other tumor types.

G) TCGA analysis of human GBM shows an immunosuppressive TME characterized by a higher ratio of PD-L1-positive cells to T cells (PD-L1:CD3), as compared to other tumor types.

H) TCGA data of human GBM as in (F) and (G), shown as a 3D projection.

### Figure S2

## A

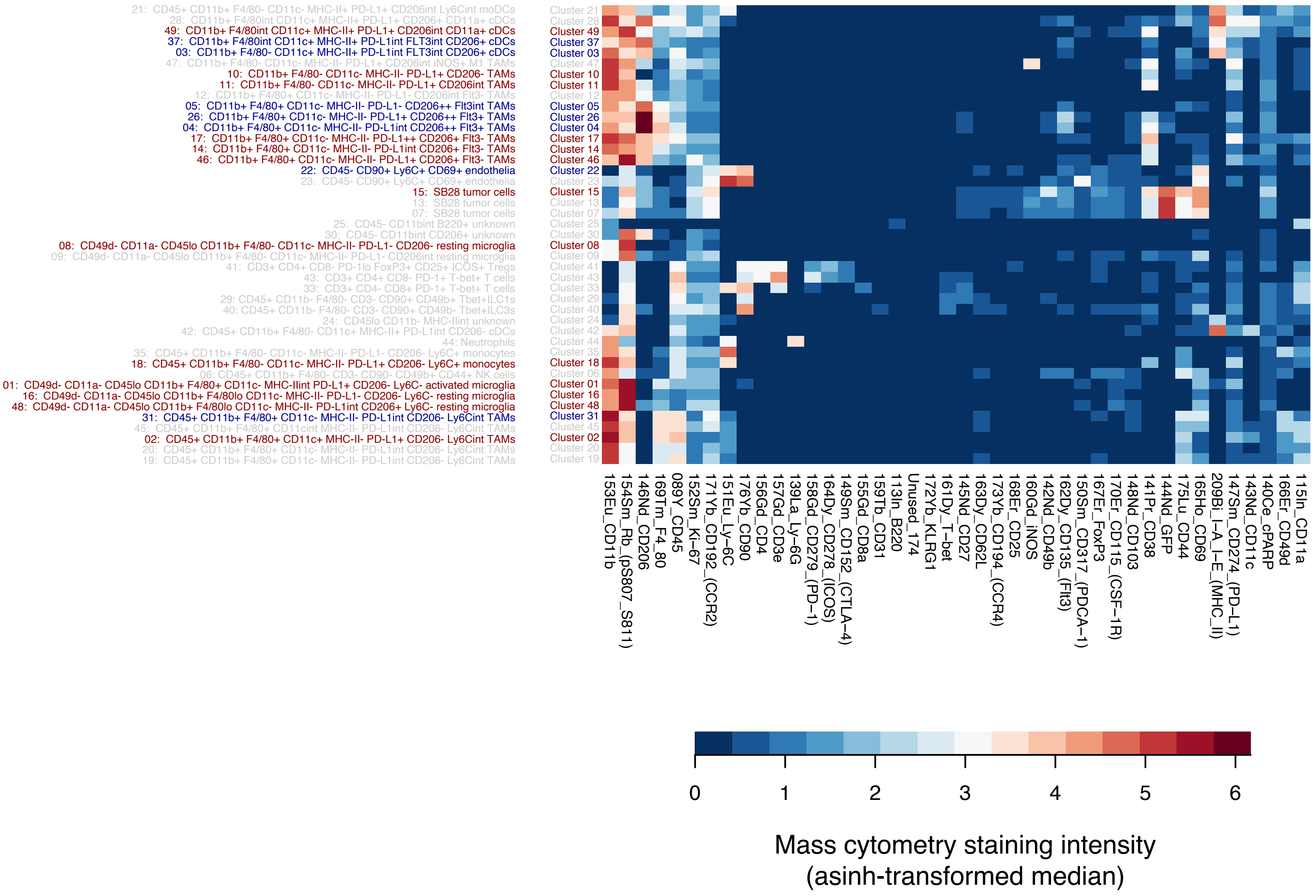
21: CD45+ CD11b+ F4/80- CD11c- MHC-II+ PD-L1+ CD206int Ly6Cint moDCs
28: CD11b+ F4/80int CD11c+ MHC-II+ PD-L1+ CD206+ CD11a+ cDCs
49: CD11b+ F4/80int CD11c+ MHC-II+ PD-L1+ CD206int CD11a+ cDCs
37: CD11b+ F4/80int CD11c+ MHC-II+ PD-L1int FLT3int CD206+ cDCs
03: CD11b+ F4/80- CD11c+ MHC-II+ PD-L1int FLT3int CD206+ cDCs
47: CD11b+ F4/80- CD11c- MHC-II- PD-L1+ CD206int iNOS+ M1 TAMs
10: CD11b+ F4/80- CD11c- MHC-II- PD-L1+ CD206- TAMs
11: CD11b+ F4/80- CD11c- MHC-II- PD-L1+ CD206int TAMs
12: CD11b+ F4/80+ CD11c- MHC-II- PD-L1- CD206int Flt3- TAMs
05: CD11b+ F4/80+ CD11c- MHC-II- PD-L1- CD206++ Flt3int TAMs
26: CD11b+ F4/80+ CD11c- MHC-II- PD-L1+ CD206++ Flt3+ TAMs
04: CD11b+ F4/80+ CD11c- MHC-II- PD-L1int CD206++ Flt3+ TAMs
17: CD11b+ F4/80+ CD11c- MHC-II- PD-L1++ CD206+ Flt3- TAMs
14: CD11b+ F4/80+ CD11c- MHC-II- PD-L1int CD206+ Flt3- TAMs
46: CD11b+ F4/80+ CD11c- MHC-II- PD-L1+ CD206+ Flt3- TAMs
22: CD45- CD90+ Ly6C+ CD69+ endothelia
23: CD45- CD90+ Ly6C+ CD69+ endothelia
15: SB28 tumor cells
13: SB28 tumor cells
07: SB28 tumor cells
25: CD45- CD11bint B220+ unknown
30: CD45- CD11bint CD206+ unknown
08: CD49d- CD11a- CD45lo CD11b+ F4/80- CD11c- MHC-II- PD-L1- CD206- resting microglia
09: CD49d- CD11a- CD45lo CD11b+ F4/80- CD11c- MHC-II- PD-L1- CD206int resting microglia
41: CD3+ CD4+ CD8- PD-1lo FoxP3+ CD25+ ICOS+ Tregs
43: CD3+ CD4+ CD8- PD-1+ T-bet+ T cells
33: CD3+ CD4- CD8+ PD-1+ T-bet+ T cells
29: CD45+ CD11b- F4/80- CD3- CD90+ CD49b+ Tbet+ILC1s
40: CD45+ CD11b- F4/80- CD3- CD90+ CD49b- Tbet+ILC3s
24: CD45lo CD11b- MHC-IIint unknown
42: CD45+ CD11b+ F4/80- CD11c+ MHC-II+ PD-L1int CD206- cDCs
44: Neutrophils
35: CD45+ CD11b+ F4/80- CD11c- MHC-II- PD-L1- CD206- Ly6C+ monocytes
18: CD45+ CD11b+ F4/80- CD11c- MHC-II- PD-L1+ CD206- Ly6C+ monocytes
06: CD45+ CD11b+ F4/80- CD3- CD90- CD49b+ CD44+ NK cells
01: CD49d- CD11a- CD45lo CD11b+ F4/80+ CD11c- MHC-IIint PD-L1+ CD206- Ly6C- activated microglia
16: CD49d- CD11a- CD45lo CD11b+ F4/80lo CD11c- MHC-II- PD-L1- CD206- Ly6C- resting microglia
48: CD49d- CD11a- CD45lo CD11b+ F4/80lo CD11c- MHC-II- PD-L1int CD206+ Ly6C- resting microglia
31: CD45+ CD11b+ F4/80+ CD11c- MHC-II- PD-L1int CD206- Ly6Cint TAMs
45: CD45+ CD11b+ F4/80+ CD11cint MHC-II- PD-L1int CD206- Ly6Cint TAMs
02: CD45+ CD11b+ F4/80+ CD11c+ MHC-II- PD-L1+ CD206- Ly6Cint TAMs
20: CD45+ CD11b+ F4/80+ CD11c- MHC-II- PD-L1int CD206- Ly6Cint TAMs
19: CD45+ CD11b+ F4/80+ CD11c- MHC-II- PD-L1int CD206- Ly6Cint TAMs
Cluster 21
Cluster 28
Cluster 49
Cluster 37
Cluster 03
Cluster 47
Cluster 10
Cluster 11
Cluster 12
Cluster 05
Cluster 26
Cluster 04
Cluster 17
Cluster 14
Cluster 46
Cluster 22
Cluster 23
Cluster 15
Cluster 13
Cluster 07
Cluster 25
Cluster 30
Cluster 08
Cluster 09
Cluster 41
Cluster 43
Cluster 33
Cluster 29
Cluster 40
Cluster 24
Cluster 42
Cluster 44
Cluster 35
Cluster 18
Cluster 06
Cluster 01
Cluster 16
Cluster 48
Cluster 31
Cluster 45
Cluster 02
Cluster 20
Cluster 19
153Eu_CD11b
154Sm_Rb_(pS807_S811)
146Nd_CD206
169Tm_F4_80
089Y_CD45
152Sm_Ki−67
171Yb_CD192_(CCR2)
151Eu_Ly−6C
176Yb_CD90
156Gd_CD4
157Gd_CD3e
139La_Ly−6G
158Gd_CD279_(PD−1)
164Dy_CD278_(ICOS)
149Sm_CD152_(CTLA−4)
155Gd_CD8a
159Tb_CD31
113In_B220
Unused_174
172Yb_KLRG1
161Dy_T−bet
145Nd_CD27
163Dy_CD62L
173Yb_CD194_(CCR4)
168Er_CD25
160Gd_iNOS
142Nd_CD49b
162Dy_CD135_(Flt3)
150Sm_CD317_(PDCA−1)
167Er_FoxP3
170Er_CD115_(CSF−1R)
148Nd_CD103
141Pr_CD38
144Nd_GFP
175Lu_CD44
165Ho_CD69
209Bi_I−A_I−E_(MHC_II)
147Sm_CD274_(PD−L1)
143Nd_CD11c
140Ce_cPARP
166Er_CD49d
115In_CD11a
0
1
2
3
4
5
6
Mass cytometry staining intensity
(asinh-transformed median)

#### Figure S2.

A) Heatmap of median marker expression in PhenoSOM populations shown in volcano plot in Figure 2D

### Figure S3

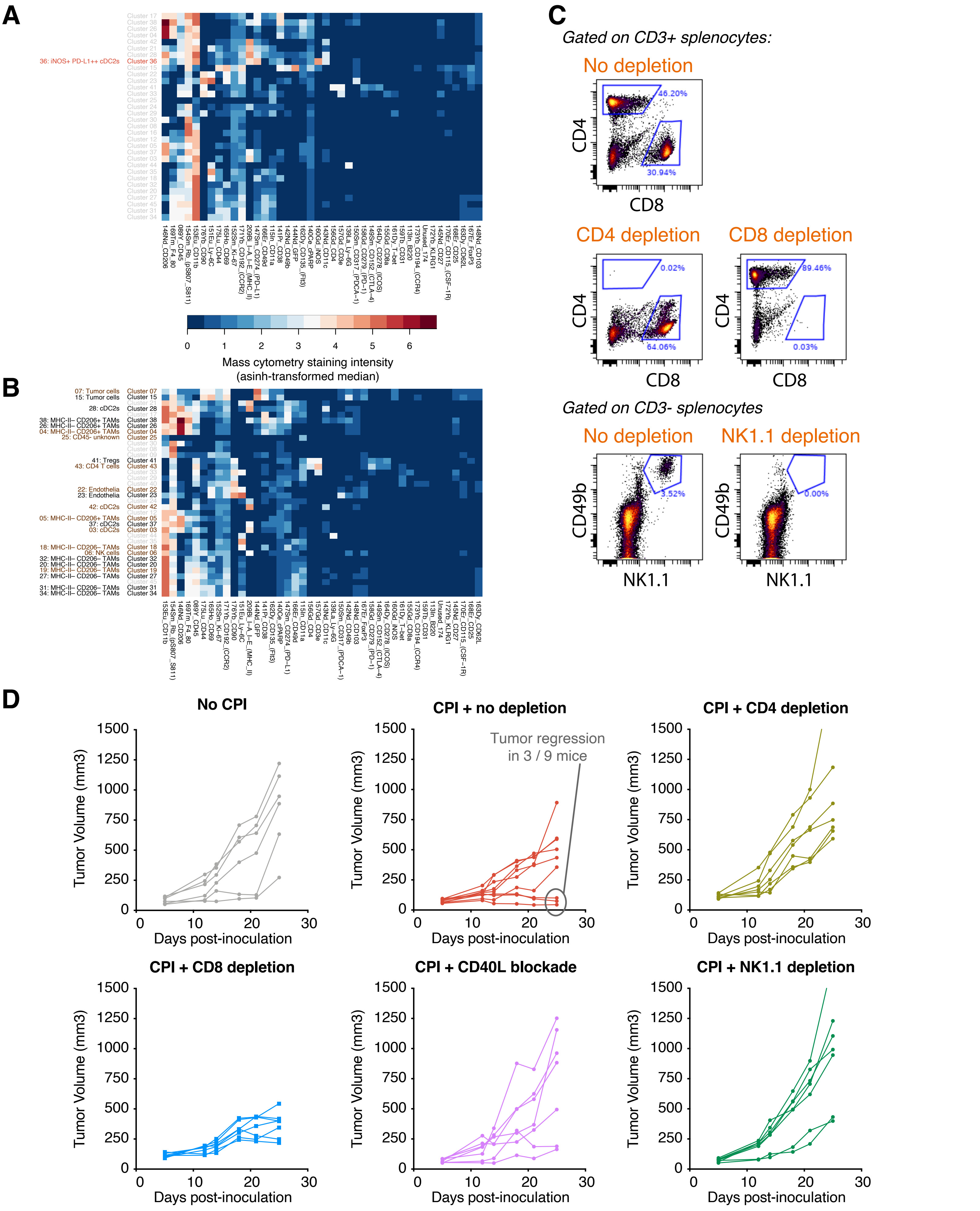
A
36: iNOS+ PD-L1++ cDC2s
Mass cytometry staining intensity
(asinh-transformed median)
B
C
Gated on CD3+ splenocytes:
No depletion
CD4
CD8
46.20%
30.94%
CD4 depletion
CD8 depletion
0.02%
64.06%
89.46%
0.03%
Gated on CD3- splenocytes
No depletion
NK1.1 depletion
CD49b
NK1.1
3.52%
0.00%
D
No CPI
CPI + no depletion
CPI + CD4 depletion
CPI + CD8 depletion
CPI + CD40L blockade
CPI + NK1.1 depletion
Tumor regression
in 3 / 9 mice
Tumor Volume (mm3)
Days post-inoculation

#### Figure S3.

A) Heatmap of median marker expression in PhenoSOM populations shown in volcano plot in Figure 3C

B) Heatmap of median marker expression in PhenoSOM populations shown in volcano plot in Figure 3D. Scale bar as in (A).

C) Flow cytometry of splenocytes from representative mice confirming successful depletion

D) Tumor growth curves of individual mice from CPI depletion cohort

### Figure S4

**A**

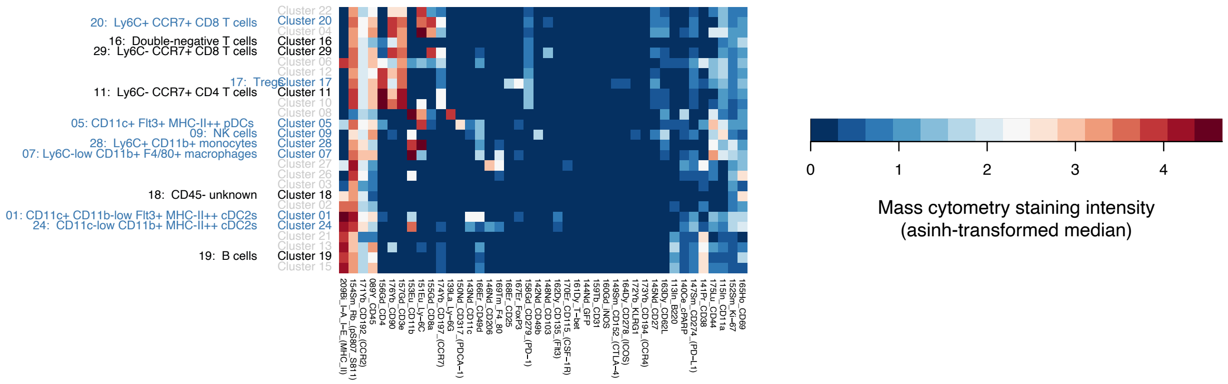

**B**

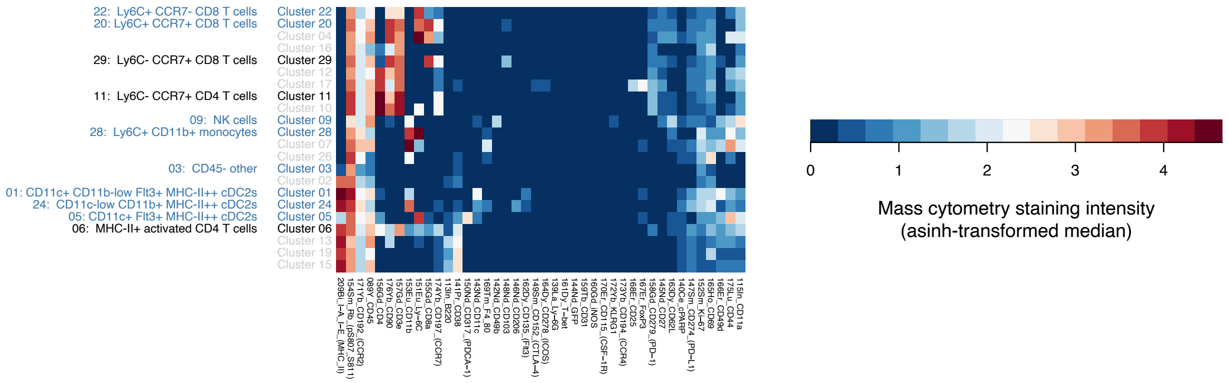

**C**

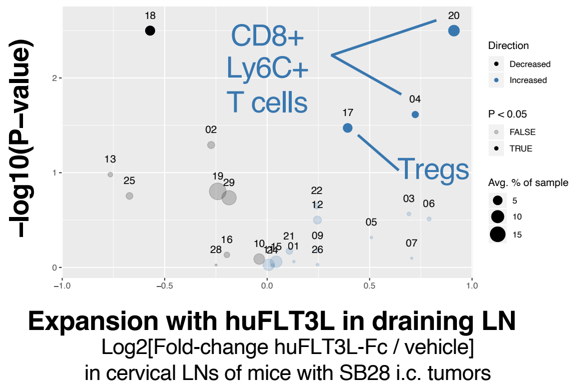

**D**

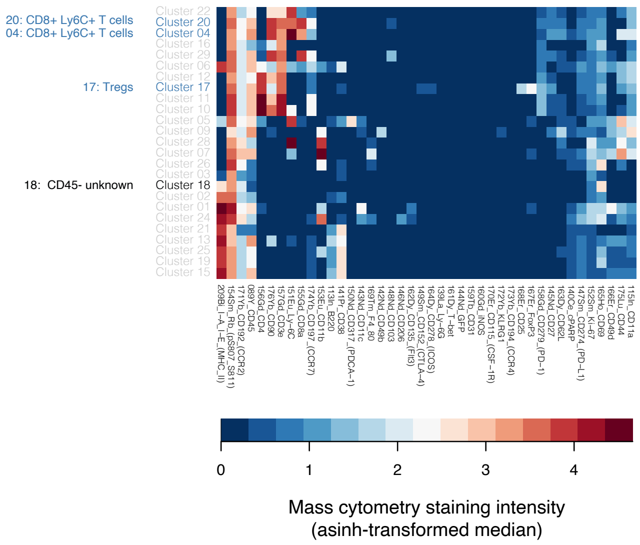

**E**

**Engraftment success rates by experiment:**

| Experiment | Site | Tumor cell line | # Injected | # Engrafted |
|---|---|---|---|---|
| 180613 | i.c. | SB28 | 14 | 14 (100%) |
| 180302 | i.c. | SB28-OVA-FL | 4 | 1 (25%) |
| 181101 | i.c. | SB28-OVA-FL | 4 | 2 (50%) |
| 180614 | s.c. | SB28 | 16 | 16 (100%) |
| 181209 | s.c. | SB28 | 44 | 44 (100%) |
| 180130 | s.c. | SB28-OVA-FL | 11 | 0 (0%) |
| 180302 | s.c. | SB28-OVA-FL | 4 | 0 (0%) |

#### Figure S4.

A) Heatmap of cluster medians in Figure 4B

B) Heatmap of cluster medians in Figure 4C

C) CyTOF shows expansion of Tregs and CD8 T cells at endpoint (Day27-33) in cervical lymph nodes of mice bearing SB28 OVA-FL i.c. tumors

D) Heatmap of marker expression in PhenoSOM clusters from Figure S4C

E) SB28-OVA-FL tumors were always rejected s.c. and frequently rejected i.c. suggesting strong immunogenicity, with limited ability to grow in the brain. Corresponds to Figure 4A.

### Figure S5

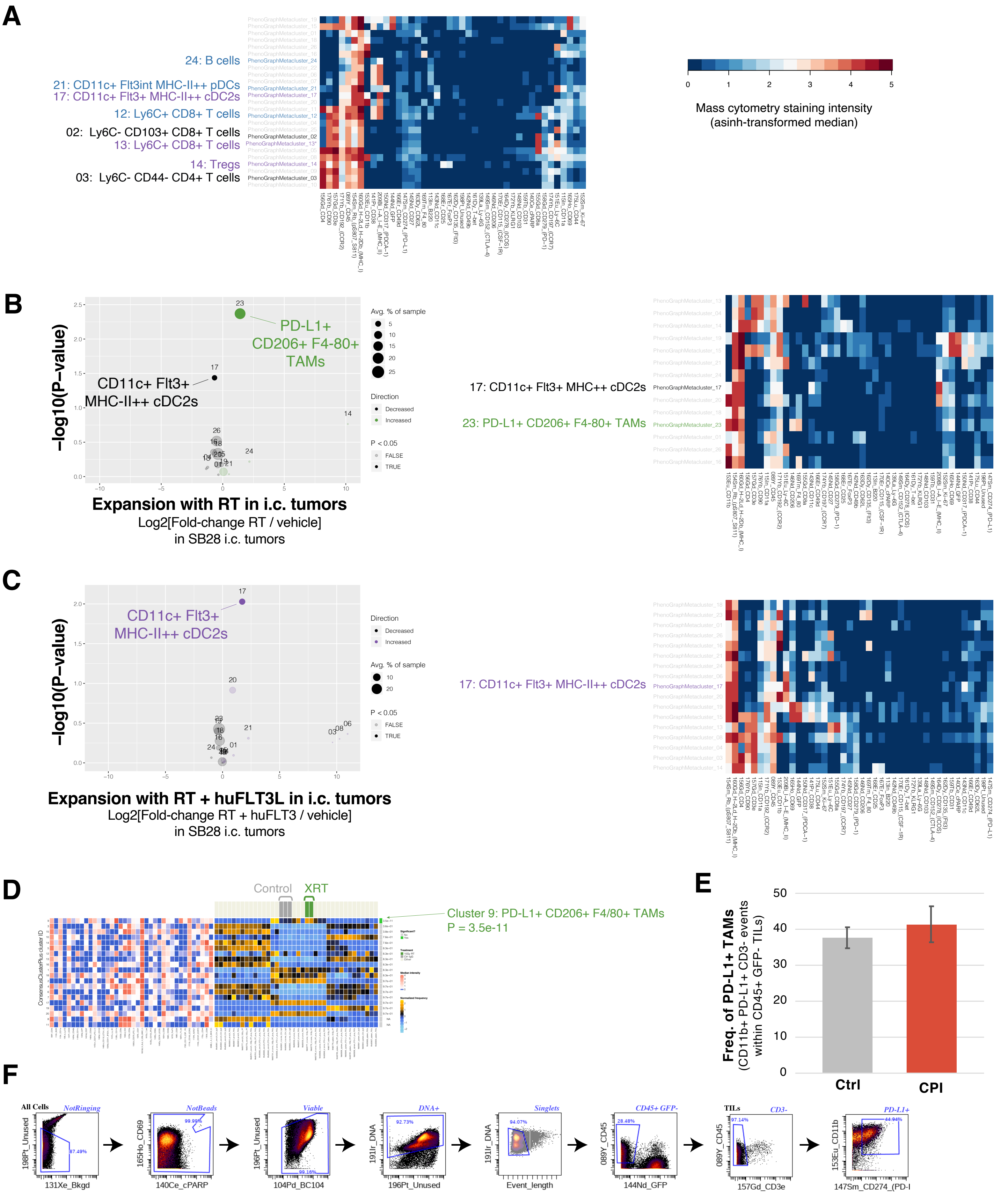
A
24: B cells
21: CD11c+ Flt3int MHC-II++ pDCs
17: CD11c+ Flt3+ MHC-II++ cDC2s
12: Ly6C+ CD8+ T cells
02: Ly6C- CD103+ CD8+ T cells
13: Ly6C+ CD8+ T cells
14: Tregs
03: Ly6C- CD44- CD4+ T cells
Mass cytometry staining intensity
(asinh-transformed median)
B
PD-L1+
CD206+ F4-80+
TAMs
CD11c+ Flt3+
MHC-II++ cDC2s
−log10(P−value)
Avg. % of sample
Direction
Decreased
Increased
P < 0.05
FALSE
TRUE
Expansion with RT in i.c. tumors
Log2[Fold-change RT / vehicle]
in SB28 i.c. tumors
17: CD11c+ Flt3+ MHC++ cDC2s
23: PD-L1+ CD206+ F4-80+ TAMs
C
CD11c+ Flt3+
MHC-II++ cDC2s
−log10(P−value)
Expansion with RT + huFLT3L in i.c. tumors
Log2[Fold-change RT + huFLT3 / vehicle]
in SB28 i.c. tumors
17: CD11c+ Flt3+ MHC-II++ cDC2s
D
Control
XRT
Cluster 9: PD-L1+ CD206+ F4/80+ TAMs
P = 3.5e-11
E
Freq. of PD-L1+ TAMs
(CD11b+ PD-L1+ CD3- events
within CD45+ GFP- TILs)
Ctrl
CPI
F
All Cells
NotRinging
NotBeads
Viable
DNA+
Singlets
CD45+ GFP-
TILs
CD3-
PD-L1+
198Pt_Unused
131Xe_Bkgd
165Ho_CD69
140Ce_cPARP
196Pt_Unused
104Pd_BC104
191Ir_DNA
196Pt_Unused
191Ir_DNA
Event_length
089Y_CD45
144Nd_GFP
089Y_CD45
157Gd_CD3e
153Eu_CD11b
147Sm_CD274_(PD-l

#### Figure S5.

A) Heatmap of marker expression in PhenoSOM clusters from Figure 5E and 5F

B) A single dose of radiation (10 Gy) induces PD-L1+ CD206+ (M2) TAMs and reduces DCs in SB28 i.c. tumors at endpoint. Volcano plot compares abundance of immune cell populations (clusters) in SB28 intracerebral tumors treated with radiation therapy (RT) (green) versus vehicle-treated (gray). Tumors were stained with the CyTOF mouse antibody panel. Statistically significant clusters in volcano plots are highlighted in opaque color and indicated with a cell type label. Diameter of the circle indicates the mean frequency of cells in the sample assigned to that cluster. Heatmap (right panel) indicates manually defined cluster phenotypes and median intensity of antibody staining in each cluster. Scale bar as in (A).

C) Radiation (1 x 10 Gy) + FLT3L restores DCs in SB28 i.c. tumors at endpoint, compared to RT alone. Volcano plot and heatmap as in (B).

D) The differential abundance analysis from (B) was repeated using the CATALYST+diffcyt analysis pipeline from Nowicka et al 2019 to compare their performance on an identical dataset (see **Online Supplemental Methods**). The Nowicka pipeline identified a cluster (Cluster 9) that is significantly enriched in RT vs. Ctrl_IgG and matches the phenotype of PhenoSOM Cluster 23 (PD-L1+ CD206+ F4-80+ TAMs) shown in (B).

E) CPI does not alter the frequency of PD-L1 TAMs in SB28 i.c. tumors at endpoint

F) Gating hierarchy for the column graph in (E)

### Figure S6

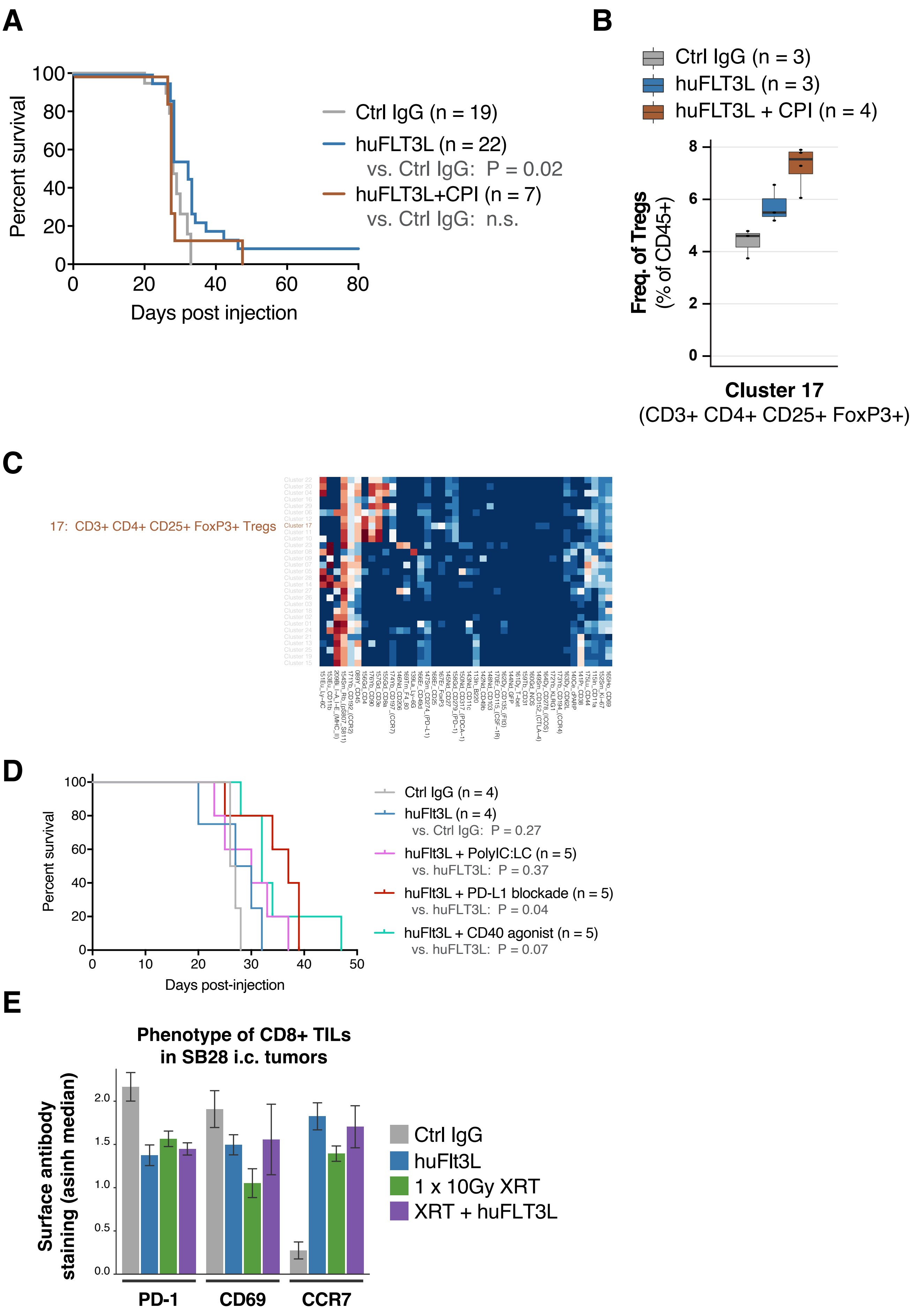
A
Percent survival
Days post injection
Ctrl IgG (n = 19)
huFLT3L (n = 22)
vs. Ctrl IgG: P = 0.02
huFLT3L+CPI (n = 7)
vs. Ctrl IgG: n.s.
B
Ctrl IgG (n = 3)
huFLT3L (n = 3)
huFLT3L + CPI (n = 4)
Freq. of Tregs
(% of CD45+)
Cluster 17
(CD3+ CD4+ CD25+ FoxP3+)
C
17: CD3+ CD4+ CD25+ FoxP3+ Tregs
D
Percent survival
Days post-injection
Ctrl IgG (n = 4)
huFlt3L (n = 4)
vs. Ctrl IgG: P = 0.27
huFlt3L + PolyIC:LC (n = 5)
vs. huFLT3L: P = 0.37
huFlt3L + PD-L1 blockade (n = 5)
vs. huFLT3L: P = 0.04
huFlt3L + CD40 agonist (n = 5)
vs. huFLT3L: P = 0.07
E
Phenotype of CD8+ TILs
in SB28 i.c. tumors
Surface antibody
staining (asinh median)
PD-1
CD69
CCR7
Ctrl IgG
huFlt3L
1 x 10Gy XRT
XRT + huFLT3L

#### Figure S6.

A) Adding αPD-1/αCTLA-4 CPI counteracts the benefit of FLT3L

B) In cervical lymph nodes at endpoint (Day 27-33), CyTOF shows that Tregs are expanded by huFLT3L alone and even more when combined with αPD-1/αCTLA-4 CPI. Box plot shows frequency of Tregs (Cluster 17: CD3+ CD4+ CD25+ FoxP3+) as a percentage of CD45-positive cells.

C) Heatmap of marker expression in Cluster 17, same dataset as (B)

D) Kaplan-Meier survival curves of signal-finding cohort to explore FLT3L combination treatments

E) Changes in the surface immunophenotypes of tumor-infiltrating T-cells in SB28 intracerebral tumors treated with FLT3L, XRT or FLT3L + XRT. Values are transformed median intensity of CyTOF measurements.
